# Characterization and diversification of AraC/XylS family regulators guided by transposon sequencing

**DOI:** 10.1101/2023.07.21.550116

**Authors:** Allison N. Pearson, Matthew R. Incha, Cindy Ho, Matthias Schmidt, Jacob B. Roberts, Alberto A. Nava, Jay D. Keasling

**Affiliations:** Joint BioEnergy Institute, 5885 Hollis Street, Emeryville, CA 94608, USA; Biological Systems & Engineering Division, Lawrence Berkeley National Laboratory, Berkeley, CA 94720, USA; Department of Plant and Microbial Biology, University of California, Berkeley, CA 94720, USA; Institute of Applied Microbiology-iAMB, Aachen Biology and Biotechnology-ABBt, RWTH Aachen University, Aachen, Germany; Joint Program in Bioengineering, University of California, Berkeley/San Francisco, CA 94720, USA; Department of Chemical and Biomolecular Engineering, University of California, Berkeley, CA 94720, USA; Institute for Quantitative Biosciences, University of California, Berkeley, CA 94720, USA; The Novo Nordisk Foundation Center for Biosustainability, Technical University of Denmark, Denmark; Center for Synthetic Biochemistry, Institute for Synthetic Biology, Shenzhen Institutes for Advanced Technologies, Shenzhen, China

## Abstract

In this study, we explored the development of engineered inducible systems. Publicly available data from previous transposon sequencing assays were used to identify regulators of metabolism in *Pseudomonas putida* KT2440. For the AraC-family regulators (AFRs) represented in this data, we posited AFR/promoter/inducer groupings. Eleven promoters were characterized for a response to their proposed inducers in *P. putida*, and the resultant data were used to create and test nine two-plasmid sensor systems in *E. coli*. Several of these were further developed into a palette of single-plasmid inducible systems. From these experiments, we observed an unreported inducer response from a previously characterized AFR, demonstrated that the addition of a *P. putida* transporter improved the sensor dynamics of an AFR in *E. coli*, and identified an uncharacterized AFR with a novel potential inducer specificity. Finally, targeted mutations in an AFR, informed by structural predictions, enabled further diversification of these inducible plasmids.

**Graphical Abstract:** 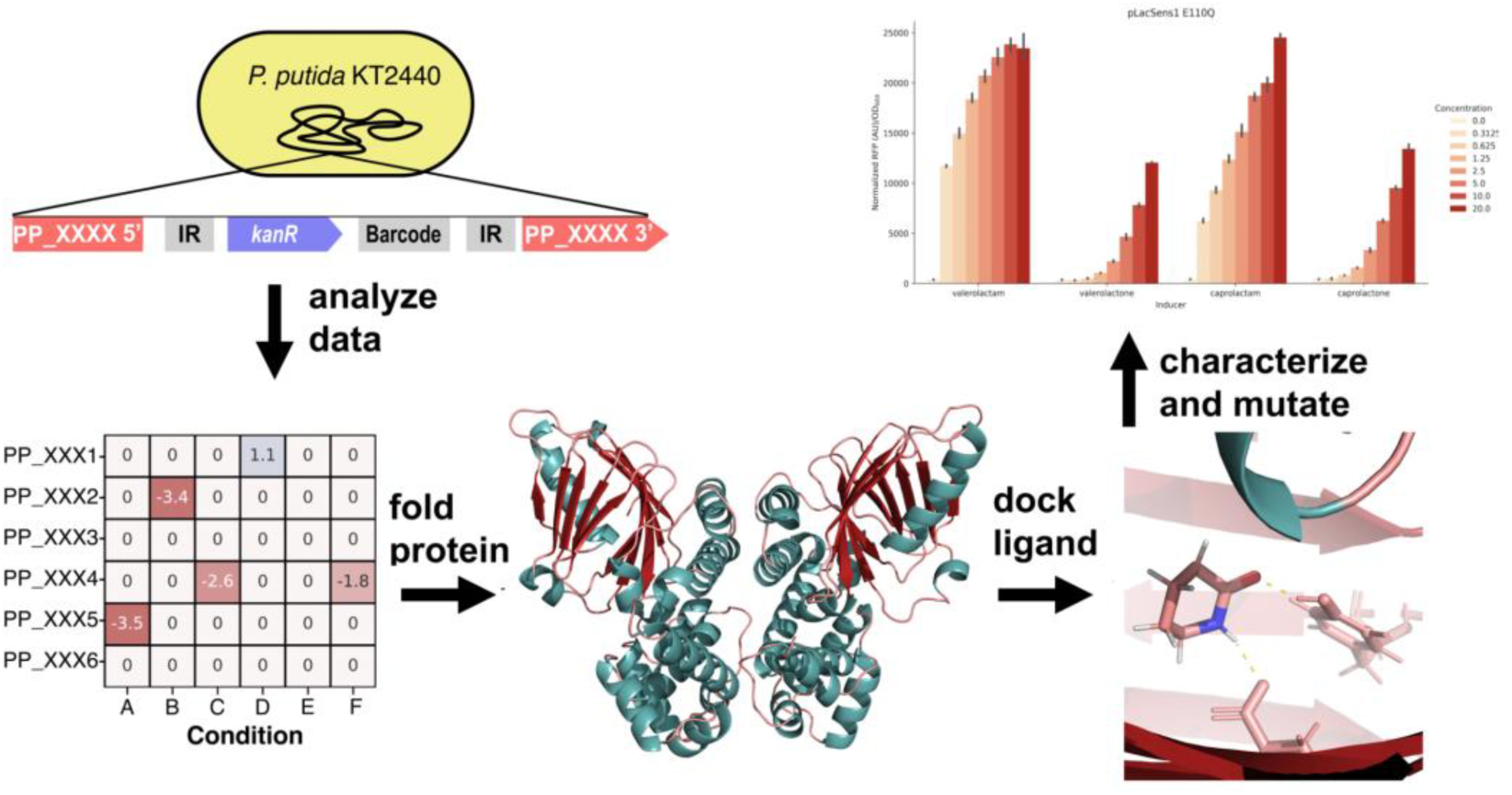

## Introduction

Rapid screening to optimize biosynthetic performance requires a method that can keep pace with process development. An allosteric transcription factor (aTF)-based biosensor can fill this need by using a fluorescent or selectable marker to quickly screen or select high-performing variants. The success of this method depends on using known aTFs with established ligands or engineering an aTF to respond to a new ligand ^1–3^. To increase probability of success, researchers should start the aTF engineering process with a well-characterized protein that has a similar ligand to the desired biochemical ^4^.

AraC/XylS-family regulators (AFRs) have shown potential as biosensors and inducible systems, due to their ability to be engineered for specific ligand recognition and previous successful applications in synthetic biology and metabolic engineering ^5–7^. AFRs are commonly found in bacteria; the most well-known and characterized are the canonical arabinose (AraC-P_BAD_) and xylose (XylS-P_xylM_) inducible systems of *Escherichia coli* and *Pseudomonas putida*, respectively, which lend the family its name ^8–10^. A repertoire of inducible AFRs would benefit bioengineering and molecular biology, each potentially offering unique benefits in terms of ligand cost or induction dynamics. Additionally, a high-throughput method for identifying these regulators would advance the development of customized inducible systems and biosensors.

Randomly barcoded transposon-site sequencing (RB-TnSeq) has proven effective in identifying key genes associated with metabolic pathways and stress responses. In our previous studies, we performed 206 RB-TnSeq assays on a mutant library of *Pseudomonas putida* KT2440, revealing significant phenotypes in over 1000 genes. By analyzing these data and drawing from database annotations, we have discovered specific functions such as substrate preferences for genes with multiple paralogs ^11–14^. In these datasets we have also identified numerous transcription factors with significant fitness phenotypes that may provide evidence for their ligand specificities and target regulons ^14^.

The advancement of new protein structure prediction tools could further accelerate development of inducible systems with more diverse ligand responses through rational mutagenesis. Previous studies have relied on known protein structures or homology models of transcription factors for rational engineering ^2^. However, AlphaFold2 and RosettaFold present a new method of protein structure prediction that can elucidate structures with high fidelity ^15, 16^. Coupling AlphaFold2 structural predictions and rational mutagenesis with RB-TnSeq methods of gene identification has yet to be described, but we believe this presents a powerful new approach for the creation of novel inducible systems.

In this study, we investigated nine AFRs represented in public RB-TnSeq data. These regulators and their promoters were characterized, and single- and two-plasmid inducible systems were created and tested in *Escherichia coli*. Through these assays, we constructed functional inducible promoters from a new uncharacterized AFR with novel ligand specificity, as well as 2 other AFRs represented in the datasets. We made targeted mutations in the binding pockets of three AFRs guided by AlphaFold structure predictions and modified their activities. This demonstrates the usefulness of functional genomics and *in silico* approaches in developing new inducible systems for controlling cell function. Many additional regulators from different families, including the LysR, GntR, GerE/LuxR, and IclR families, also have significant fitness changes in publicly available RB-TnSeq datasets ^17–20^. We believe future work could employ these other transcription factors represented in RB-TnSeq datasets to generate more diverse inducible systems.

## Results

### Phenotypic identification of AraC family regulators

*Pseudomonas putida* KT2440 carries 40 distinct AraC/XylS family regulators (AFRs) in its genome, based on a search of the genome for proteins containing the conserved AFR helix-turn-helix domain, Pfam HTH_18 (PF12833) ^21^. Some have known or predicted functions; however, others lack a specific annotation. Through analyzing barcode transposon abundance sequencing (RB-TnSeq) data, we found 16 AFRs with significant (|t-score|>4, |fitness|>1) phenotypes (Figure 1)^14, 22, 23^. The regulator OplR (PP_3516) was previously identified via proteomics and later was shown to have a significant phenotype in a nitrogen-source RB-TnSeq dataset^11–14^. Other AFRs present in the data, such as gbdR (PP_0298), cdhR (PP_0305), benR (PP_3159), pobR (PP_3538), and argR (PP_4482), either have predicted functions based on homology or have been previously characterized and align with our fitness data ^24–28^. Based on prior literature, co-fitness, and/or proximity to other metabolic genes, we assigned regulon predictions for 9 of the 16 AFRs in the data (Table 1).

**Figure 1.**
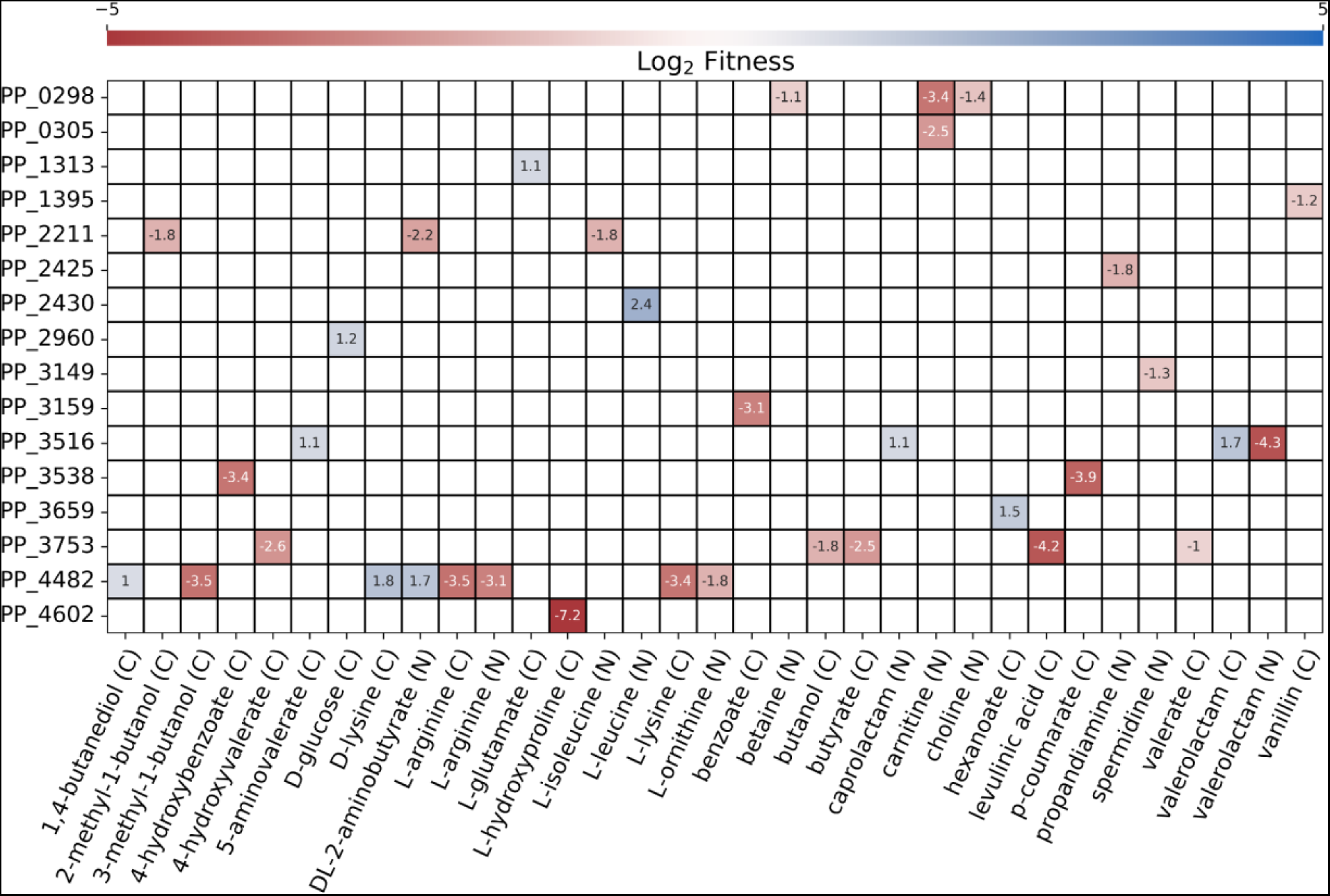
Fitness data for AraC family transcription regulators with significant fitness changes (|t-score| > 4, |fitness| >1). Fitness changes that did not meet the cutoff are not shown.

**Table 1.** Promoters, predicted transcriptional regulators, and predicted in-vivo ligands for each system tested in this study, along with the fold induction between the maximum measured normalized RFP values and the uninduced controls. n=3, error = standard deviation.

| Predicted AFR | Predicted target promoter | Predicted *in vivo* ligands | Tested ligands | Fold induction of promoters in *P. putida* |
| --- | --- | --- | --- | --- |
| PP_0298 (gdbR) | PPP_0296 | betaine | betaine | 2.2 ± 0.26 |
| PP_0305 (cdhR) | PPP_0304 | carnitine | carnitine | 270 ± 13 |
| PP_3149 | PPP_3148 | spermidine | spermidine | 67 ± 8.5 |
| PP_3538 (pobR) | PPP_3537 | 4-hydroxybenzoate | 4-hydroxybenzoate | 160 ± 31 |
| PP_3753 | PPP_3754 | 3-hydroxybutyrate, or 3-hydroxyvalerate | 3-hydroxybutyrate<br>levulinic acid | 1.3 ± 0.32<br>6.9 ± 0.65 |
| PP_4602 | PPP_1259 | L-hydroxyproline | L-hydroxyproline | 310 ± 41 |
| PP_3159 (benR) | PPP_3161 | benzoate | benzoate | 38 ± 5.2 |
| PP_3159 (benR) | PPP_3160 | benzoate | benzoate | 1.7 ± 0.20 |
| PP_4482 | PPP_4486 | L-arginine | L-arginine | 6.5 ± 0.24 |
| PP_4482 | P<sub>PP_4481</sub> | L-arginine | L-arginine | 11 ± 2.1 |
| PP_2211 | P<sub>PP_2213</sub> | 2-methylbutyrate, or<br>2-methylbutyryl-CoA | 2-methylbutyrate<br>2-methylbutanol | 52 ± 4.7<br>19 ± 0.62 |

### Promoter characterization in P. putida

From the predicted regulon for the AFRs, we extracted the 200 base pairs upstream of the genes and constructed reporter plasmids with the promoters driving red fluorescent protein (RFP) expression. We then tested these reporter plasmids (plasmids pIP12-pIP23) in *P. putida* (strains sIP1-sIP12) for their response to the predicted ligand in minimal medium (Figure 2A) (Tables S1, S2). The predicted 3-hydroxybutyrate (P_PP_3754_) and betaine (P_PP_0296_) responsive promoters (sIP7, sIP1) both showed minimal induction in the corresponding test conditions. However, P_PP_3754_ (sIP7) did show a weak (6.9-fold induction) response to levulinic acid, which agrees with our RB-TnSeq data, indicating that the AFR/promoter pair PP_3753/P_PP_3754_ might be involved in the regulation of levulinic acid or 3-hydroxypentanoyl-CoA catabolism. Interestingly, with maximum inductions of 6.5 and 11-fold, the two L-arginine responsive promoters (sIP8, sIP9) had similar dose dependent responses while sharing little homology in their nucleotide sequence. With 67-fold maximum induction at an inducer concentration of 5 mM, P_PP_3148_ (sIP3) exhibited a moderate response to spermidine; however, the cells failed to reach a high OD at spermidine concentrations higher than 10 mM, likely due to toxicity from spermidine itself or a downstream metabolite. Similarly, the promoter P_PP_3161_ (sIP4) was induced 38-fold above background at 10 mM benzoate, but was hampered at 20 mM benzoate due to a presumed toxic effect on cell growth. Both 2-methylbutyrate and 2-methylbutanol induced moderate expression from the predicted promoter P_PP_2213_, (sIP11) at 52 and 19-fold induction, respectively. P_PP_3537_ (sIP6) demonstrated a strong response to 4-hydroxybenzoate, with a maximum of 160-fold induction following supplementation with p-HB. Finally, the predicted carnitine (P_PP_0304_) and L-hydroxyproline (L-HPro) (P_PP_1259_) responsive promoters (sIP2 and sIP12, respectively) both demonstrated high expression in the presence of their respective inducers and tight repression, with 270 and 310 -fold induction (Table 1).

**Figure 2.**
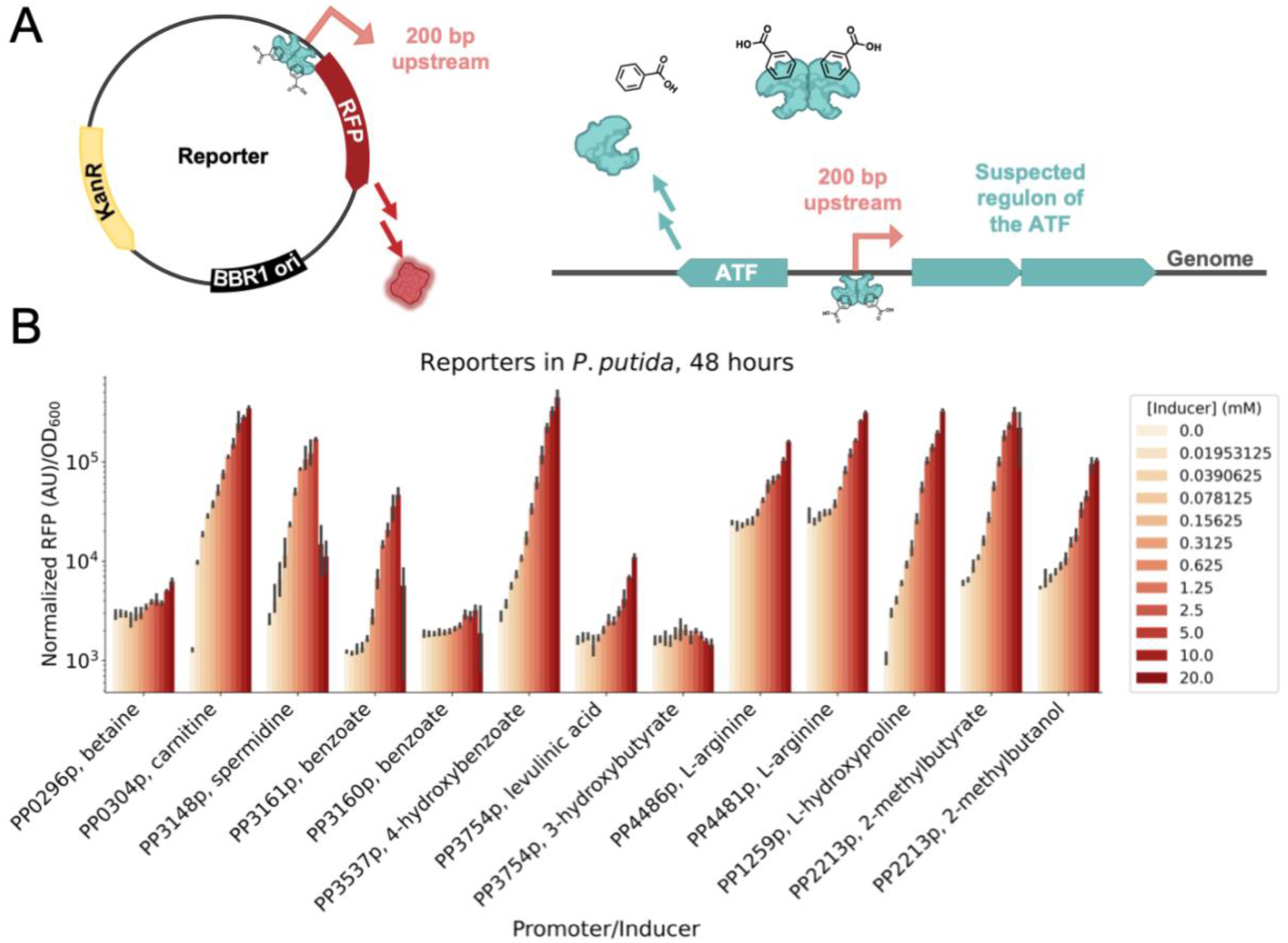
A) Schematic of the reporter plasmids used in *P. putida* to assay for inducibility. B) Barchart showing the induction of the promoter (x-axis) with their cognate inducer in *P. putida* grown in minimal medium supplemented with 10 mM glucose. Promoters (200 bp upstream of start codon) for each gene were induced with the predicted ligand. P_PP_0296_ (sIP1) with betaine, P_PP_0304_ (sIP2) with carnitine, P_PP_3148_ (sIP3) with spermidine, P_PP_3161_ (sIP4) and P_PP_3160_ sIP5) with benzoate, P_PP_3537_ (sIP6) with para-hydroxybenzoate, P_PP_3754_ (sIP7) with 3-hydroxybutyrate and levulinic acid, P_PP_4486_ (sIP8) and P_PP_4481_ (sIP9) with L-arginine, P_PP_1259_ (sIP12) with L-hydroxyproline, and P_PP_2213_ (sIP11) with 2-methylbutyrate and 2-methylbutanol. (n=3, error bars=95% confidence interval).

### Development of inducible systems in *E. coli*

Following the promoter assay in *P. putida*, we next sought to validate the requirement for the corresponding AFRs via heterologous expression in *E. coli.* We used a previously described two-plasmid strategy, cloning the AFRs into arabinose-inducible ‘regulator’ vectors (plasmids pIP1-pIP9) and transforming them into *E. coli* carrying their cognate reporter plasmids (plasmids pIP12-pIP23) – the same constructs used to test the inducible activity of the promoters in *P. putida* (Figure 3A, Table S1) ^23^. The resultant strains (strains sIP13-sIP24) allowed us to vary both the AFR expression level and inducer concentration in a high-throughput manner (Table S2).

We constructed a two-plasmid system using BenR as a test case. BenR is an AFR with a well-characterized response to benzoate in regulating the 69 base-pair *Pb* promoter ^29^. In the two-plasmid system (sIP16), BenR is regulated by the canonical AraC-P_BAD_ system (pIP4), and the region 200 bp upstream of PP_3161 (P_PP_3161_) was used as the promoter for RFP (pIP15) (Figure 3A). With the knowledge that this region contained the characterized *Pb* promoter, we expected induction of RFP to coincide with increasing concentrations of arabinose and benzoate ^26^. Our results indicate that the promoter region was intact and that the expression of the regulator is titratable with arabinose (Figure 3B).

We tested the eight other inducible systems in the same manner as BenR. PP_4482/P_PP_4481_ (sIP21) and PP_4482/P_PP_4486_ (sIP20) had increased RFP expression with increased expression of the regulator. However, the induction of RFP did not seem to be affected by the addition of exogenous L-arginine (Figures S2I, S2J). Similarly, PP_2211/P_PP_2213_ (sIP23) demonstrated a RFP response when the regulator was induced, but this response also did not increase when the putative inducers, 2-methylbutanol and 2-methylbutyrate, were added to the medium (Figure S2K, S2L). Likewise, PP_3538/P_PP_3537_ (sIP18) demonstrated a slight increase in RFP expression upon induction of the regulator, but there was no titratable response to the addition of 4-hydroxybenzoate, although this may be due to lack of transport across the membrane (Figures S2F, S3). There was no clear correlation with inducer or regulator expression to RFP expression in the cases of PP_0298/P_PP_0296_ (sIP13), PP_0305/P_PP_0304_ (sIP14), PP_3148/P_PP_3149_ (sIP15), or PP_3754/P_PP_3753_ (sIP19) (Figure S2A, S2B, S2C, S2D, S2E, S2G). There did appear to be a response to both inducer and regulator expression in the instance of PP_4602/P_PP_1259_ (sIP24), and this is further described in a later section.

**Figure 3.**
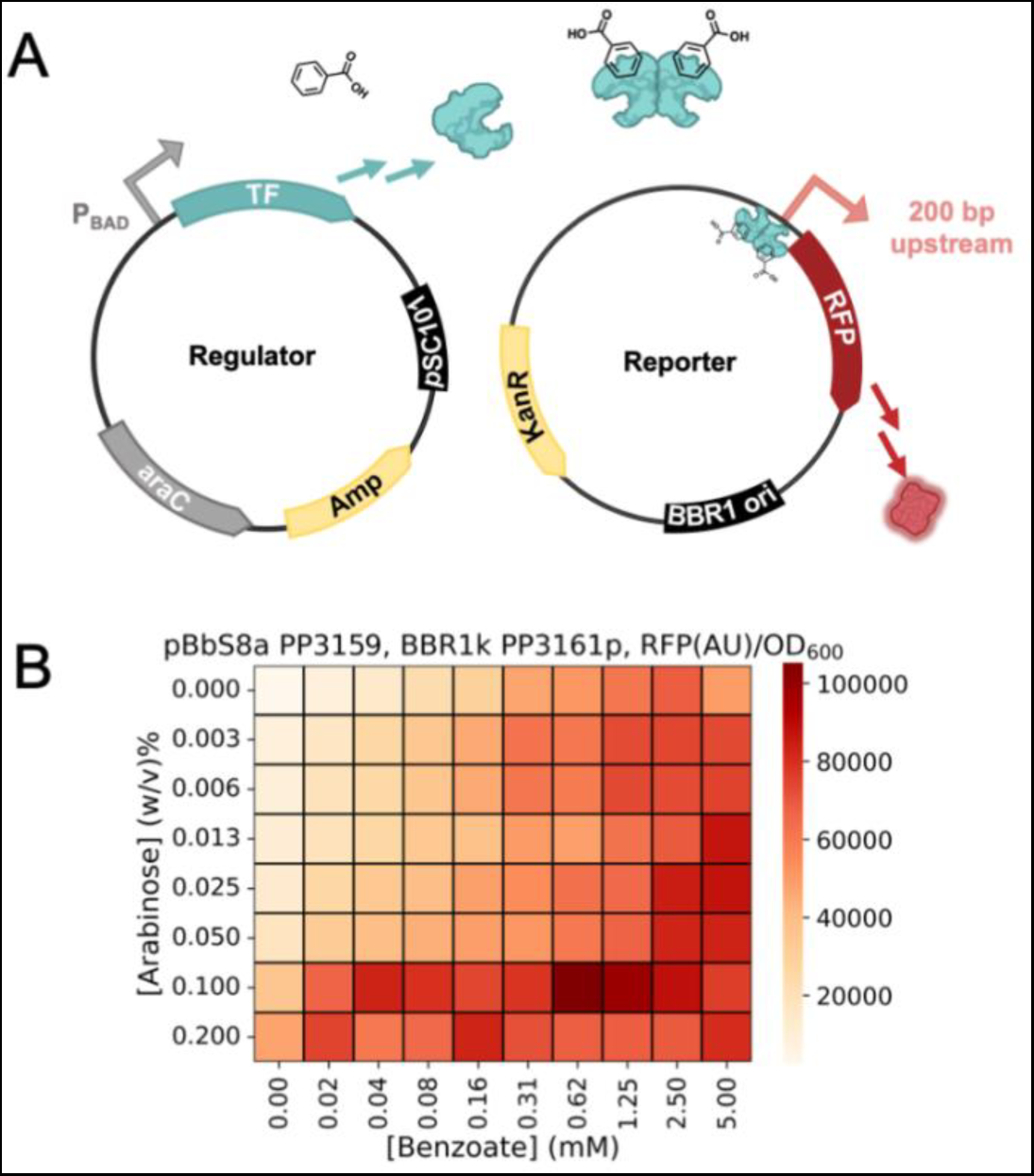
A) Schematic of two-plasmid system used in *E. coli* to assay promoter-transcription factor relationships. The transcription factor is induced on a low copy pSC101 plasmid with the addition of arabinose, and the expression of RFP is driven by the cognate promoter for the transcription factor B) Data from the two-plasmid system (sIP16) carrying the PP_3161 promoter and inducible BenR (PP_3159), n = 3. Plot of SD shown in Figure S1. Cells were cultured in LB medium for 24 hours.

### PP_3159 (BenR) responds to multiple benzoates

Following the success with the two-plasmid system, we developed three one-plasmid systems (plasmids pIP33-pIP35) employing the benzoate-inducible BenR and also tested them in *E. coli* (strains sIP39-sIP41)(Table S1, Table S2). Each one-plasmid system had a different strength constitutive promoter driving the expression of *benR* and the 200 bp upstream of PP_3161 controlling expression of RFP ^30^. Each variant showed different sensor dynamics with the p1 variant (sIP39) having the tightest repression and 38 ± 3.9 fold induction, the p3 variant (sIP41) showing the highest sensitivity and 8.7 ± 0.98 -fold induction, and the p2 variant (sIP40) showing an intermediate level of sensitivity and repression with 33 ± 6.2 -fold induction (Figures 4A, 4B).

BenR has previously been described as a highly specific benzoate-responsive transcription factor ^29, 31^, but we found that it also responds to structurally similar compounds. We tested three functionalized benzoates with our one-plasmid system containing the strong p3 promoter expressing *benR* (sIP41). Surprisingly, we found that our one-plasmid system responded to 3-methylbenzoate and salicylate in addition to benzoate (Figure 4C). The maximal induction by 3-methylbutyrate was ∼ 80% of induction by benzoate, while the maximal induction by salicylate and benzoate were roughly the same. However, the system’s response to benzoate was stronger than salicylate at low concentrations. This could be due to the inclusion of a secondary BenR binding site in our construct or because of high BenR expression from the p3 promoter, which is apparent in the high background fluorescence of this system (Figures 4A, 4B).

**Figure 4.**
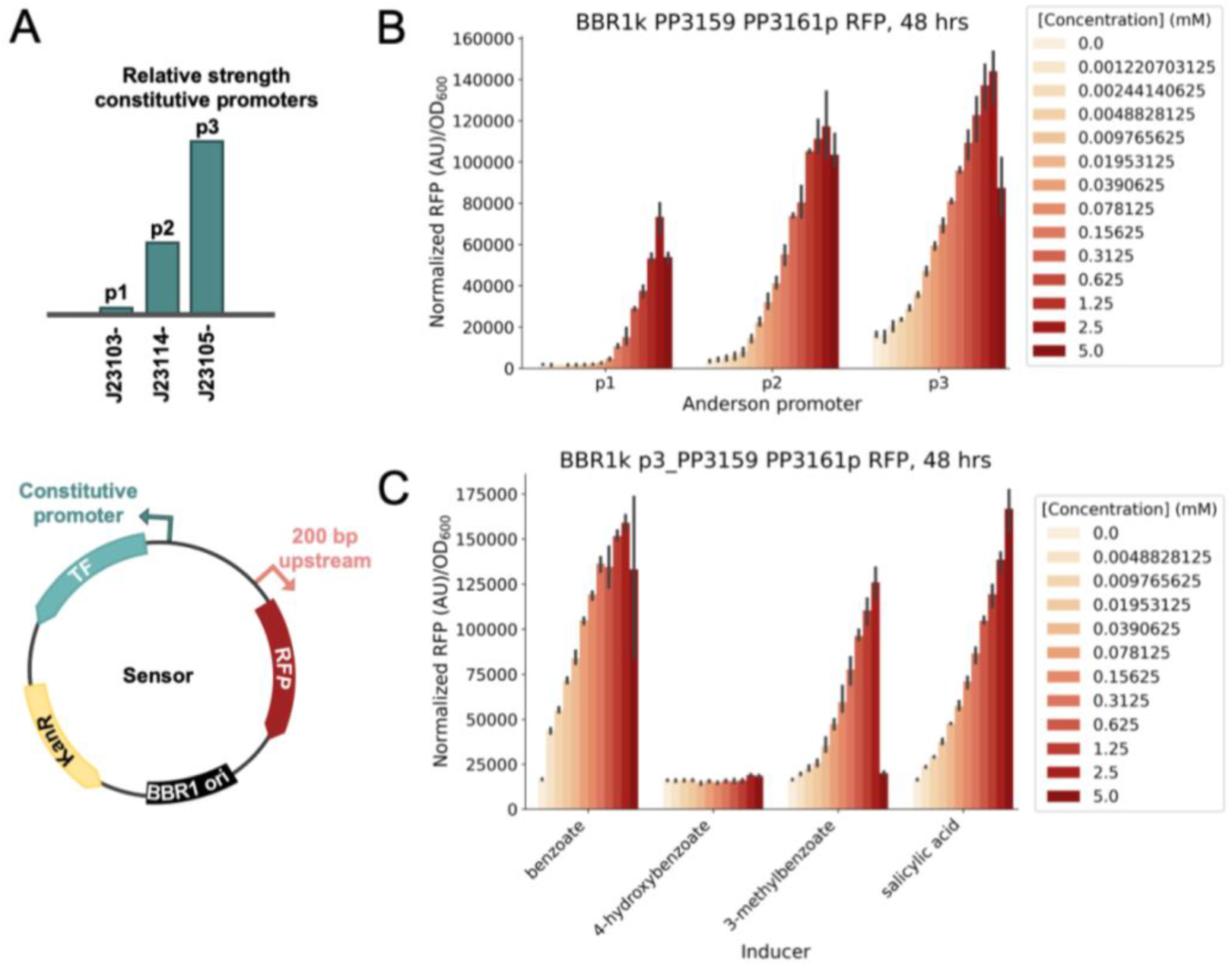
Single plasmid system for BenR (n=3, error bars=95% confidence interval). A) Schematic of the one-plasmid systems with constitutive promoters driving expression of the transcription factor, and RFP under the control of the transcription factor. The bars (not to scale) are labeled with the relative strength of the three Anderson promoters used. B) Dose dependent response bar plots showing increasing benzoate concentrations induce expression of RFP. C) Induction of RFP from the p3 single plasmid BenR system (sIP41) with functionalized benzoates. Induction studies were conducted for 48 hours in LB medium.

### PP_4602 allosteric response is enhanced with coexpression of a transporter

The AFR PP_4602 exhibited a specific phenotype (|t-score| > 4, |fitness| >1) in the RB-TnSeq data with trans-L-4-hydroxyproline (L-HPro) as the carbon source. The only other genes displaying specific phenotypes in this condition are the predicted L-HPro assimilation genes located at loci PP_1255, PP_1256, PP_1257, PP_1258, and PP_1259. A unique characteristic of PP_4602 is that it lacks a typical AFR ligand binding domain, instead it contains a N-terminal Per-Arnt-Sim (PAS) domain. This domain, as observed in other proteins, is involved in ligand binding ^32, 33^. Homology and fitness data suggest that this AFR is expected to behave similarly to LhpR (PA1261) from *Pseudomonas aeruginosa*, considering the significant sequence similarity (57% identity) of their ligand binding domains ^34^.

In *P. putida*, P_PP_1259_ (sIP12) demonstrated a pronounced response to L-HPro, which makes it a promising candidate for the creation of an inducible system in *E. coli* (Figure 2B). The PP_4602/P_PP_1259_ two-plasmid system (sIP24) exhibited a tunable response with the induction of the transcriptional regulator and the inclusion of the suitable inducer, L-HPro (Figure 5A, S4). Subsequently, we built three single-plasmid systems (plasmids pIP24-pIP26) and evaluated their response to L-HPro in *E. coli* (strains sIP30-sIP32) (Tables S1, S2). Although these inducible systems exhibited a clear response at an average of 7.1 ± 1.1 fold induction, they did not attain maximal induction at the same concentration observed in *P. putida* (Figures 2B, 5B). After adding the predicted transporter, PP_1259, to be co-transcribed with the ATF in the the single-plasmid system (pIP27-pIP29, sIP33-sIP35), maximal induction was reached at much lower L-HPro concentrations (390 µM), the average fold induction across the three systems increased to 9.8 ± 2.6, and the systems conformed to the Hill equation (Figure 5C).

**Figure 5.**
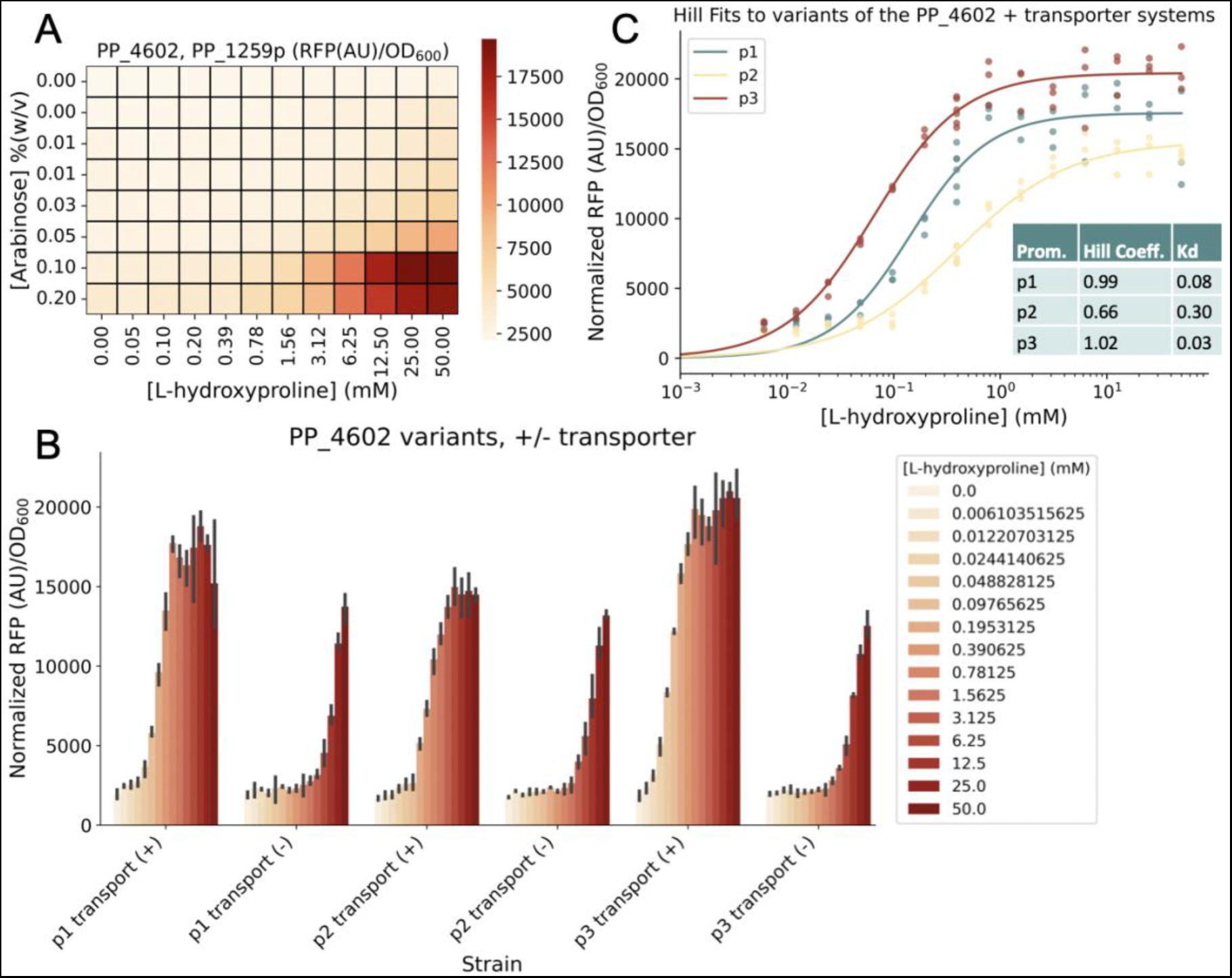
Systems for PP_4602 (LhpR). A) Data from the two-plasmid system carrying the PP_1259 promoter and inducible lhpR (sIP24), n = 3. Plot of SD shown in Figure S4. B) Barplot showing the inducibility of the PP_4602 one-plasmid systems (sIP30-sIP35) with and without the L-hydroxyproline transporter in a synthetic operon with lhpR (n=3, error bars=95% confidence interval). C) Hill function fit to the PP_4602 promoter variants with the L-hydroxyproline transporter co-expressed with PP_4602 (sIP33-sIP35). Cells were cultured in LB medium for 24 hours.

### PP_2211 putatively responds to 2-methylbutyryl-CoA

The AFR PP_2211 and the neighboring putative beta-oxidation system encoded by PP_2213-PP_2217 have significant fitness defects (|t-score| > 4, fitness < 1) in the presence of 2-methylbutanol, L-isoleucine, and DL-2-aminobutyrate, indicating that these genes are essential for the utilization of these compounds (Figure 1).^14^ In *P. putida*, all three of these compounds may share 2-methylbutyryl-CoA as a catabolic intermediate. Previous research has proposed that 2-methylbutanol is oxidized to 2-methylbutyrate, which subsequently undergoes beta-oxidation via the PP_2213-PP_2217 operon ^12^. It has also been posited that 2-aminobutyrate is catabolized through deamination to 2-oxobutyrate, which is then funneled into L-isoleucine biosynthesis and subsequent catabolism^14^. The acyl thioester 2-methylbutyryl-CoA is a known intermediate during the catabolism of L-isoleucine.^35^

This suggests that PP_2211 responds to free 2-methylbutyrate, 2-methylbutyryl-CoA, or a downstream metabolite. If the AFR PP_2211 responds to free 2-methylbutyrate, then the fitness data would suggest that some thioesterase acts upon the 2-methylbutyryl-CoA produced during growth on isoleucine and 2-aminobutyrate. It is also possible that PP_2211 may respond to 2-methylbutyryl-CoA itself or a downstream metabolite. To further examine this, we grew *P. putida* WT and ΔPP_2211 carrying the P_PP_2213_ reporter plasmid (sIP11, sIP27) with L-isoleucine, 2-methylbutyrate, and succinate as carbon sources in minimal medium (Figure 6A, Table S2). There was very little RFP expression in the ΔPP_2211 strain (sIP27), confirming that PP_2211 regulates expression from P_PP_2213_. The WT strain (sIP11) showed no significant RFP expression when strains were grown on succinate, moderate RFP expression when grown on L-isoleucine, 2-methylbutyrate, and isoleucine + succinate, and high RFP expression when grown on 2-methylbutyrate + succinate. If PP_2211 responded to free 2-methylbutyrate, we would expect that the RFP expression levels would correlate directly to the amount of 2-methylbutyrate provided, but instead we see a dependence on succinate.

PP_2211 may form a positive feedback loop with PP_2213, which encodes a CoA ligase that also has a strong fitness phenotype and has been proposed to act on 2-methylbutyrate during 2-methylbutanol metabolism ^12^. It is possible that this operon is a positive feedback loop, similar to the LacI system in *E. coli* ^36^. PP_2211 may respond to small amounts of a downstream product, such as 2-methylbutyryl-CoA, and upregulate expression of the PP_2213-PP_2217 operon. To test this hypothesis, we created a ΔPP_2213 strain and compared the response of the reporter plasmid to 2-methylbutyrate in this background (sIP28) versus the wild-type background (sIP11). Maximal RFP expression was ∼70% lower in the ΔPP_2213 background (sIP28), indicating that PP_2211 is likely responding to a downstream metabolite such as 2-methylbutyryl-CoA (Figure 6B). PP_2216 is an acyl-CoA dehydrogenase posited to act on 2-methylbutyryl-CoA, and when we tested the reporter in a ΔPP_2216 strain (sIP29), we saw a ∼70% increase in maximal RFP response (Figure 6B) ^12^. This strongly suggests that PP_2211 detects 2-methylbutyryl-CoA itself.

Finally, we sought to further probe this system in *E. coli* via a two-plasmid system. We created a strain (sIP25) that carried both a single-plasmid sensor variant (pIP30) and a plasmid with the CoA-ligase PP_2213 (pIP10) under the control of the P_BAD_ system (Figure 6D, Tables S1,S2). We varied the abundance of both the metabolite and the CoA-ligase PP_2213 by growing this strain (sIP25) with different concentrations of 2-methylbutyrate and arabinose. We found that when expression of the CoA-ligase PP_2213 was increased via the addition of arabinose, RFP expression from the sensor increased up to 2.9 ± 0.51 -fold (Figure 6E). There was no RFP expression above background when no 2-methylbutyrate was added; however, beyond the lowest concentration of 2-methylbutyrate added (9.8 μM), adding more 2-methylbutyrate did not increase the sensor response. Production of 2-methylbutyryl-CoA could be dependent on the amount of enzyme and not the amount of substrate due to feedback inhibition. 2-Methylbutyryl-CoA may allosterically inhibit the CoA ligase, similar to the *Rhodopseudomonas palustris* benzoate-CoA ligase ^37^. Together, these data indicate that PP_2211 responds to 2-methylbutyryl-CoA. We propose this AFR be named the 2-methylbutyrate regulator, TmbR.

**Figure 6.**
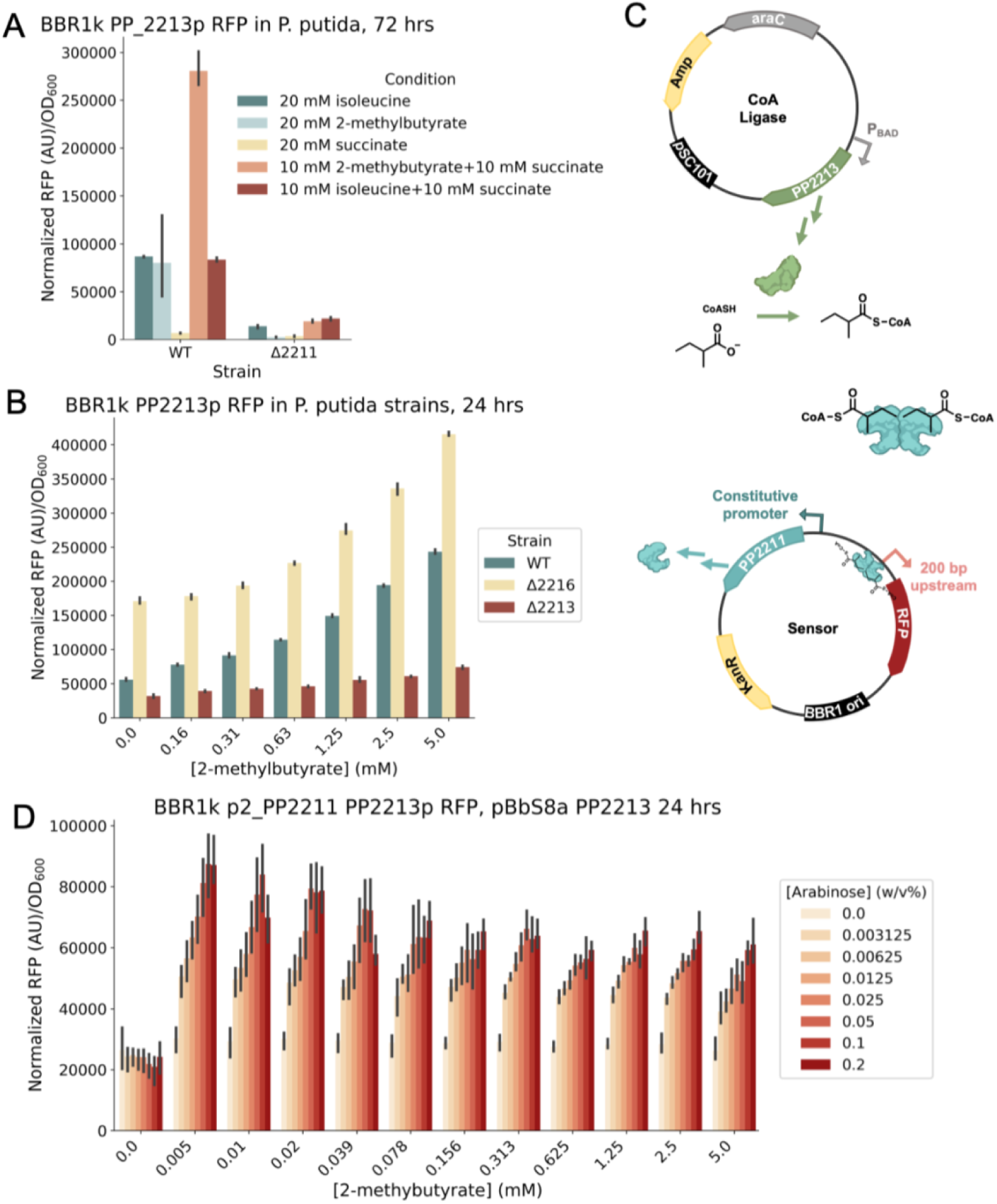
Investigation of the PP_2211/P_PP_2213_ system (n=3, error bars=95% confidence interval). A) Induction of RFP expression from P_PP_2213_ in *P. putida* WT (sIP11) and ΔPP_2211 (sIP27) background. Cultures were grown in MOPS minimal medium for 72 hours with the indicated carbon sources. B) Induction of RFP expression from P_PP_2213_ in *P. putida* under varied concentrations of 2-methylbutyrate in the *P. putida* WT (sIP11), ΔPP_2216 (sIP29), and ΔPP_2213 (sIP28) backgrounds. Cultures were grown in LB medium for 24 hours. C) Schematic of a two plasmid system used to characterize the P_PP_2213_ in *E. coli*. The CoA ligase plasmid features PP_2213 expressed by the AraC-P_BAD_ system (pIP10), while the sensor plasmid contains the PP_2211/P_PP_2213_ system (pIP30). D) RFP expression from the system shown in part C. Cultures were grown in EZ Rich defined medium for 24 hours, supplemented with varying concentrations of arabinose and 2-methylbutyrate.

### Rational mutations for altered ligand specificities

Following development of the inducible systems for *E. coli*, we sought to alter the substrate specificity of the regulators via targeted mutagenesis. Using our local implementation of AlphaFold2, LBL Foldy, we predicted the protein structures for the AFRs investigated in this work ^16, 38–40^ (Table S3). We noted PP_3516 (OplR), PP_4602 (LhpR), and PP_3159 (BenR) contained pockets larger than a water molecule in their predicted ligand binding domains. Using *in silico* docking either with Autodock Vina or SwissDock ^41, 42^, we then docked the ligands to the structures (Figures 7, S5A, S6A). Following these predictions we generated rational mutations in PP_4602, BenR, and OplR to alter their ligand specificity.

The AFR PP_4602 was annotated as having a non-canonical start site TTG, and the naive AlphaFold structure prediction shows that the first 20 amino acids encode a disordered, low predicted accuracy alpha helix. Upon further interrogation, we identified an internal methionine with a canonical ATG start codon 20 amino acids into the annotated sequence. We chose this as the start site for the rest of our *in silico* analyses. The signal sensing domain of PP_4602 is smaller than the other AFRs in this study while still containing a distinct cavity within it, but this cavity was not large enough to accommodate L-HPro. Following *in silico* repacking via sidechain and backbone energy minimization in FoldIt-Standalone, the pocket could accommodate L-HPro, and ligand binding was predicted with SwissDock (Table S3) ^41–44^.

With this new prediction, we identified four residues with possible H-bond interactions with the ligand. In one predicted binding mode, K46 and K123 appear to be H-bonding with the carboxylic acid group of L-HPro. R62 was within H-bonding distance to the hydroxyl, and Y52 was near the secondary amine (Figure S5A). With this information, we predicted a mutation in K46 or K123 into an H-bond acceptor like glutamate or aspartate could confer a response to S-or R-4-hydroxy-2-pyrrolidinone (S or R-HPyr). However, following individual mutations at K46 and K123 (plasmids pIP40-pIP41), the sensor did not respond to S/R-HPyr or L-HPro in *E. coli* (strains sIP46-sIP47) (Figure S5B, TableS1-S2).

We hypothesized mutations in BenR could also enable a response to benzyl-alcohol and benzaldehyde. H32 appeared to form pi-stacking while Y61 and Y115 made H-bonds with the benzoate ligand docked to the structure (Figure S6A). We mutated residues H32A, Y61F, and Y115F (pIP37-pIP39) in an attempt to yield a transcription activation response to benzyl alcohol and benzaldehyde in *E. coli* (sIP43-sIP45). These mutations also abolished all inducible activity (Figure S6B and C). However, our characterization of both the LhpR and BenR mutants indicated that the residues we chose may be essential for signal transduction or ligand binding.

**Figure 7.**
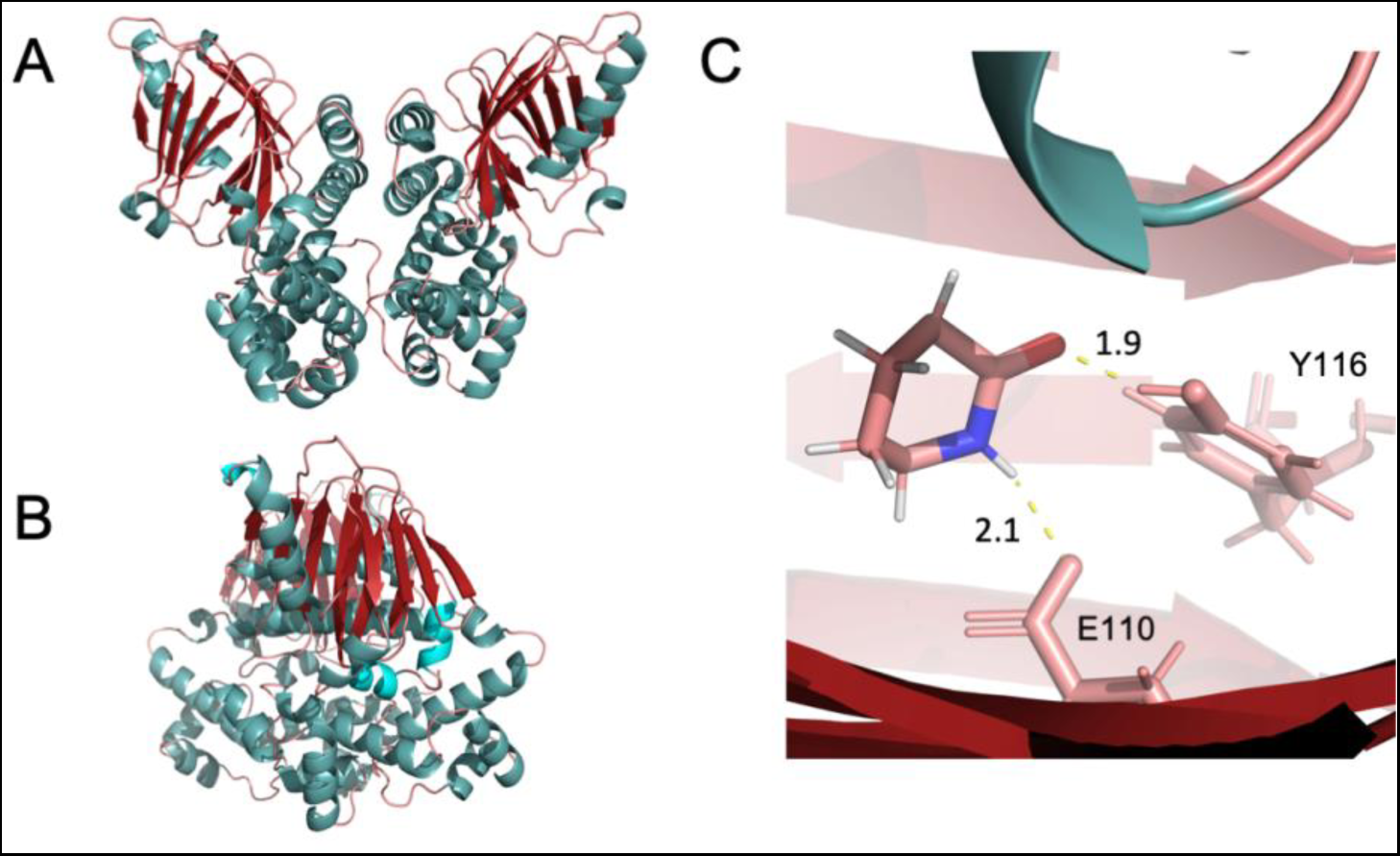
AlphaFold-predicted structure of OplR. A) ‘Front’ view of OplR structure. B) OplR structure rotated 90 degrees along the vertical axis. C) Zoomed in view of OplR ligand binding site docked with valerolactam via AutoDock Vina showing E110 and Y116 with hydrogen bonds depicted to the valerolactam ligand. Distances are in angstroms.

Finally, we chose the caprolactam and valerolactam responsive ATF, OplR, as our final test case for mutational analysis, and successfully changed its substrate specificity ^23^. We identified the residues defining the ligand binding pocket of OplR and found six residues with polar groups. It appeared that Y116 and E110 could form H-bonds with the oxygen and nitrogen of the amide, respectively. We hypothesized that a mutation at E110 to an H-bond donor such as glutamine would confer a response to valerolactone and caprolactone. E110Q (pIP43, sIP49) resulted in inducible expression by both valero- and capro-lactam and lactone (Figure 8, Table 2). Surprisingly, we also found the E110Q mutation enabled a response to many more cyclic ligands (Figure 8, Table 2). Furthermore, an alanine-substituted mutant, E110A (pIP42, sIP48), was more selective in ligand specificity, favoring cyclohexanone and ethyl-valerolactam. OplR E110A (sIP48) also exhibited approximately 60% lower background fluorescence than wildtype and 70% lower background than E110Q (sIP49), while exhibiting the highest -fold induction (33 ± 1.2, for ethyl-valerolactam) out of the tested OplR variants (Figure 8, Table 2).

**Figure 8.**
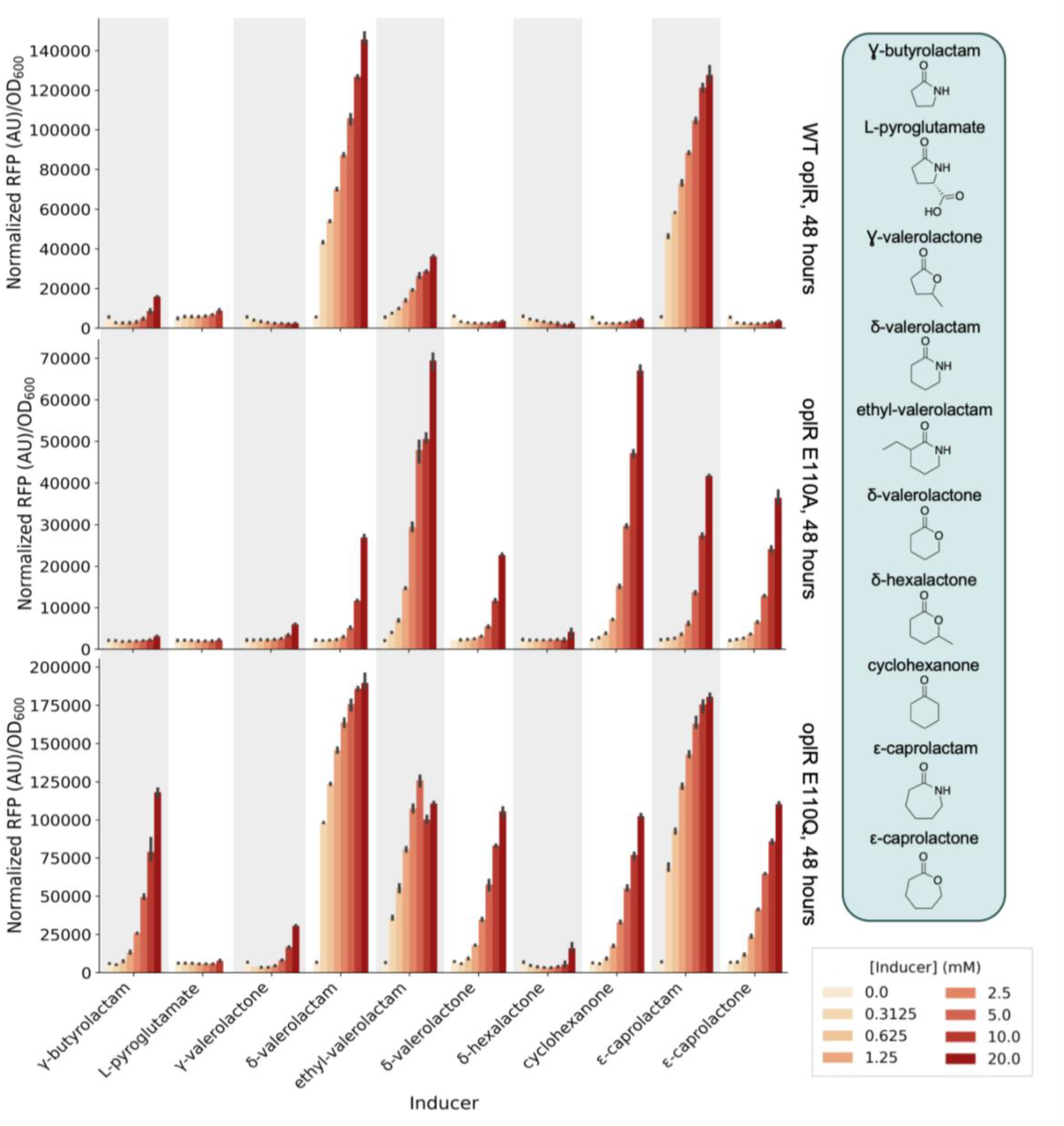
Response of OplR wild-type and mutant variants (sIP48-sIP49) in *E. coli* to ten cyclic inducers: γ-butyrolactam, L-pyroglutamate, γ-valerolactone, δ-valerolactam, ethyl-valerolactam, δ-valerolactone, δ-hexalactone, cyclohexanone, ε-caprolactam, and ε-caprolactone. The mutants and WT demonstrate different substrate specificities. Cultures were grown in LB medium with each inducer for 48 hours (n=3, error bars=95% confidence interval).

**Table 2.**
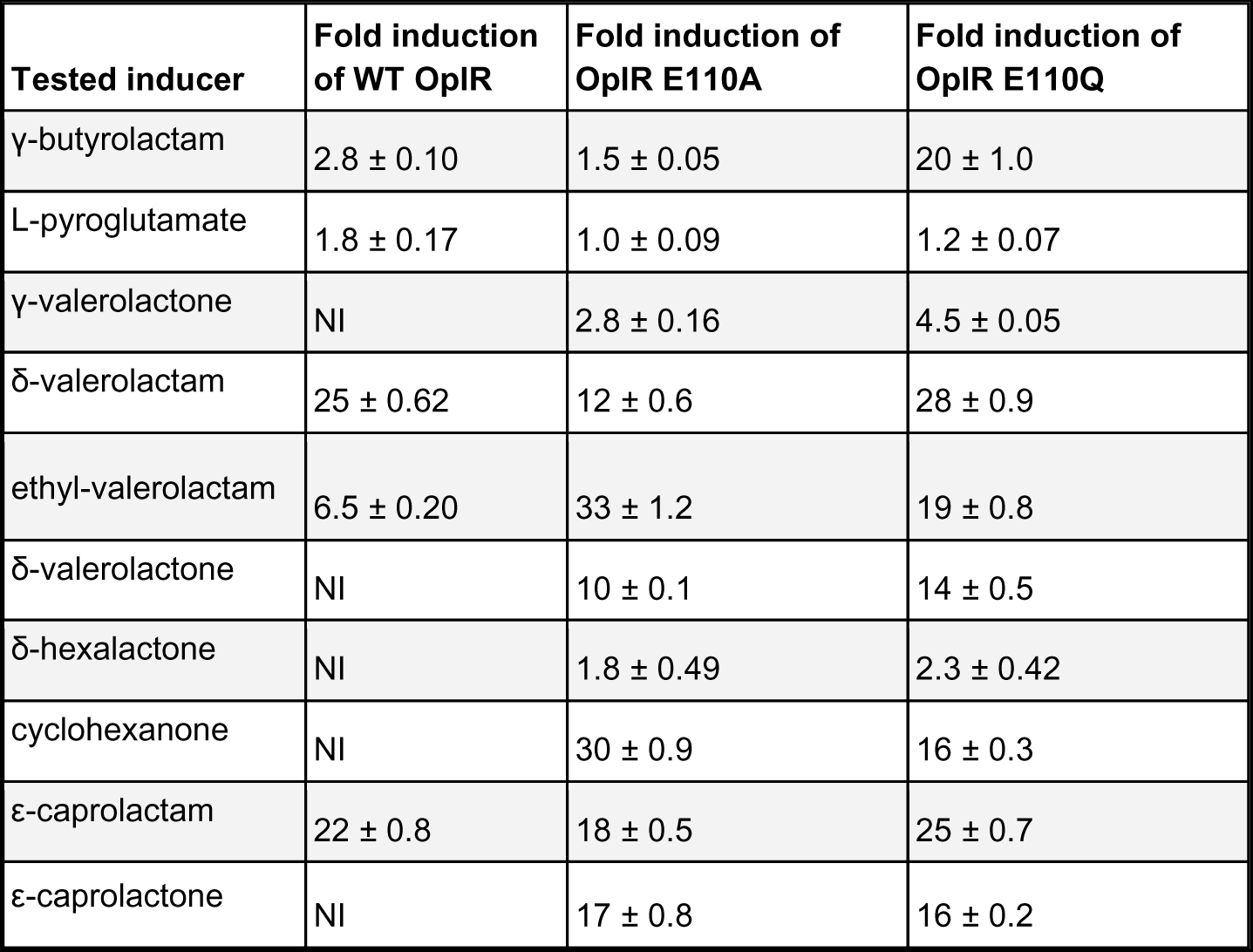
OplR WT and mutants with their response to tested lactam and lactone inducers. The fold induction values shown are between the maximum measured normalized RFP values and the uninduced controls. Fold induction values <1 are indicated by NI (no induction). n = 3, error = standard deviation.

## Discussion

Here we have described a process for identifying and characterizing transcription factors using public data derived from the high throughput functional genomics method, RB-TnSeq, and its application to the AraC-family transcription factors in *P. putida*. We first employed RB-TnSeq data to hypothesize AFR-promoter-inducer pairings. Then we reintroduced these promoter regions into the native host (*Pseudomonas putida* KT2440) as reporter systems (sIP1-sIP12) and observed whether they enabled titratable gene expression in the presence of the suspected inducer. With functional knockouts in downstream genes, this series of reporters could be employed in *P. putida* for bioengineering or general microbiology studies. We next developed a series of inducible plasmids (pIP24-pIP35) that enable expression with diverse inducers in *E. coli* (sIP30-sIP41). These systems can further be employed in a manner akin to the canonical P_BAD_ inducible system. Finally, we demonstrated how protein structure predictions via AlphaFold2 and FoldIt can be further used to gain a deeper understanding of these proteins through targeted mutations either resulting in disrupted induction activity or in modulated activity, further expanding the diversity of these regulators.

Interestingly, PP_4602 shows homology to the regulator of the L-HPro degradation system in *P. aeruginosa*, LhpR, but it is distal from the other genes involved in L-HPro degradation ^34^. This regulator and its promoter are located near identical predicted transposable elements. This may indicate that the regulator migrated via a spurious transposition or recombination event. The inducible systems (pIP24-pIP26) developed from this regulator show both low background in rich medium and high induction in the presence of L-HPro when tested in *E. coli* (sIP30-sIP32). Incorporating the L-HPro transporter from *P. putida* into the one-plasmid systems (pIP27-pIP29) resulted in peak induction at lower concentrations of HPro in *E. coli* (sIP33-sIP35) (Figure 5A). However, in all tested constructs, the use of constitutive promoters with different strengths to drive expression of the ATF did not appear to have a significant effect on sensor dynamics. Our Alphafold2 structure predictions indicated that the true start codon might be 60 bp downstream of the currently annotated start codon, which could explain this observation.

The benzoate catabolism regulator, BenR (PP_3159) is proximal to a hypothetical protein, PP_3160. After testing this gene’s promoter in *P. putida*, we showed that it is not responsive to benzoate, unlike the promoter region 5’ of PP_3161 (*Pb*). The exact function of PP_3160 is still unknown. The three BenR-derived single-plasmid systems (pIP33-pIP45) we developed have different sensitivities and dynamic ranges in *E. coli* (sIP39-sIP41) and could be employed as expression systems for metabolic engineering applications. We found that our p3 single-plasmid construct (sIP41) was strongly induced by 3-methylbenzoate and salicylate, in addition to benzoate. This differs from previously described reports that BenR demonstrates little to no response to these molecules ^31^. We did use a longer sequence as the ‘promoter’ region than in previous work; our 200 base-pair ‘promoter’ for RFP expression contains the canonical 69 base-pair *Pb* promoter as well as a portion of the upstream gene, PP_3160, and it is possible that our constructs included a secondary BenR binding site ^26^. Previous work has found that adding an additional BenR binding site to the *Pb* promoter enabled a weak response to 3-methylbenzoate.^29^ Another explanation could be that our constructs yielded higher intracellular concentrations of BenR than are present natively, amplifying the effect of weak binding to these 3-methylbenzoate and salicylate. The use of constitutive promoters driving expression of the transcription factor may be useful in identifying weak activities to inducers and guide protein engineering efforts.

From 2-methylbutanol, L-isoleucine, and 2-aminobutyrate RB-TnSeq data, we identified an AFR (PP_2211, TmbR) with a potentially novel ligand specificity for 2-methylbutyryl-CoA. This would be a novel ligand specificity for AFRs, as no prior AFR has been described to have a response to an acyl-CoA, and TmbR may be applicable as a biosensor for pathways relying on these molecules. While we could not identify a discreet ligand binding pocket for this protein based on AlphaFold structural predictions, directed evolution methods could be conducted to enhance the response to 2-methylbutyryl-CoA or enable a response to another acyl-CoA of interest.

Finally, we successfully leveraged an AlphaFold structure to make informed mutations and change the substrate specificity of the characterized valerolactam responsive transcription factor, OplR ^16, 38, 45^. AutoDock Vina resolved a potential binding mode for valerolactam in the predicted ligand binding domain of OplR ^39, 40^. A single mutation, E110A, in this site enabled broader substrate specificity and responses to ligands which had no inducible activity in the wildtype protein and reduced the inducible response to valerolactam (Figure 8). We hypothesized that another mutation, E110Q, would enable an allosteric response to lactones. Not only did the E110Q mutant (sIP49) respond to valerolactone and caprolactone, it also responded to cyclohexanone, enhanced the response to ethyl-valerolactam and butyrolactam, and provided a weak response to gamma-valerolactone (Table 2). These constructs could prove to be a starting point for further mutation of OplR and diversification of its inducer space. Lactams, lactones, and their derivatives are targets for bioproduction, and increasing the range of these compounds that genetically-encoded biosensors can detect may aid in future engineering efforts ^46, 47^.

The identification of these regulators enabled the rapid development of transcription factor-based inducible systems that can be used in all fields requiring engineered protein/RNA expression. The systems derived from the benzoate and L-HPro responsive ATFs/promoters (PP_3159/P_PP3161_, PP_4602/P_PP1259_) were functional in both *P. putida* and *E. coli*. Experiments in both organisms also indicated that PP_2211 (TmbR) likely responds to 2-methylbutyryl-CoA, a novel inducer type for an ATF. Finally, we created targeted mutations in the AFR OplR that altered its inducer specificity.

We have explored the AraC family of transcription factors in *P. putida* using public RB-TnSeq data, reporter assays, inducible systems, and AlphaFold2. The approach utilized in this work could be used to study additional transcription factor families in *P. putida* or other organisms. Such characterization of transcription factors expands our fundamental biological knowledge and our tools for metabolic engineering and synthetic biology.

## Methods

### Plasmids and Strains

The bacterial strains and plasmids utilized in this research are detailed in Tables S1 and S2. Any strains and plasmids generated during this research are accessible via the public JBEI registry. (https://public-registry.jbei.org/folders/792). We designed all plasmids through the Device Editor and Vector Editor software, and j5 software was used for designing all primers that were involved in the plasmid construction process ^48–50^. We assembled the plasmids through Gibson Assembly following standard procedures ^51^. Routine isolation of the plasmids was performed using the Qiaprep Spin Miniprep kit from Qiagen, USA, while all primers were procured from Integrated DNA Technologies (IDT, Coralville, IA). DNA sequencing was conducted through Primordium (Monrovia, CA)

### Chemicals, media, growth conditions

*E. coli* and *P. putida* were cultivated in Lysogeny Broth (LB) Miller medium (sourced from BD Biosciences, USA) at a temperature of 37°C and 30°C, respectively. When necessary, strains were also cultured in EZ-Rich medium (obtained from Teknova, Hollister, CA) with a 10 mM glucose supplement. Minimal media experiments were conducted with MOPs buffered minimal medium prepared according to LaBauve et al ^52^. As needed, the cultures were enriched with kanamycin (50 mg/L from Sigma-Aldrich, USA), carbenicillin (100 mg/L from Sigma-Aldrich, USA) or gentamicin (30 mg/L from Sigma-Aldrich, USA). All additional chemicals were acquired from Sigma-Aldrich (Sigma-Aldrich, USA).

### Fluorescence assays

Endpoint assays were carried out in 96-deep well plates (procured from Corning Costar, model 3960). Each well was filled with 500 μL of a medium that included necessary ligands, antibiotics, and/or inducers, with an inoculation of 1% v/v sourced from overnight cultures. These plates were sealed with an AeraSeal film (from Excel Scientific, model AC1201-02) and incubated at 30°C or 37°C on a 250 rpm shaker rack. After 24 and/or 48 hours where specified, 100 μL from every well was distributed into a black, clear-bottom 96-well plate (Corning Costar, 3603) to measure optical density and fluorescence using an Biotek Synergy H1 (Agilent, Santa Clara, CA) plate reader. Optical density was evaluated at 600 nm (OD_600_), and fluorescence was assessed using an excitation wavelength of 535 nm, an emission wavelength of 620 nm, and a manually set gain of 100.

### Structure predictions and docking

Structure predictions were conducted with the Foldy implementation of AlphaFold2 ^38, 45^. All AFRs were folded as monomers and dimers and were docked to the predicted ligand based on the RB-TnSeq datasets. Docking was conducted in the Foldy UI using AutoDock Vina as the docking algorithm ^39, 40^. For LhpR (PP_4602), the protein structure prediction for the dimer was repacked using Foldit-Standalone, and docking was conducted using SwissDock ^41, 43, 44^. No boundaries for ligand binding were specified in any of the docking experiments. Folds for all AFRs and dimers are available as public structures in the LBL implementation of Foldy, at https://foldy.lbl.gov/ with the tag “AraC” (Table S1).

### *Gene knockouts in* Pseudomonas putida

Gene knockouts in *Pseudomonas putida* were made as previously described ^12^. The allelic exchange vector pMQ30 was used for homologous recombination with *sacB* counterselection. Homology arms of roughly 1000 bp each, including the start and stop codons, were cloned into the pMQ30 vector. These vectors were then electroporated into *E. coli* S17 and subsequently conjugated into *P. putida*. Transconjugants were selected for on Pseudomonas isolation agar (Difco) supplemented with 30 mg/mL gentamicin. Putative knockouts were selected for on LB plates with no NaCl and 10% w/v sucrose and screened via PCR with primers flanking the target gene.

### RB-TnSeq datasets and AFR identification

RB-TnSeq data were collected from the fitness browser (fit.genomics.lbl.gov). These data included experiments from numerous publications, including data from the original paper describing the library ^11, 12, 14, 22, 53, 54^. AFRs were identified from the datasets by setting a t-score cutoff of ±4 and screening for proteins containing the Pfam HTH_18 (AraC family helix-turn-helix) domain ^21^.

## Acknowledgements

We thank Bridget Luckie and Peter Winegar for providing feedback on figure designs, and Michael Cronce, Isaac Donnell, and Aidan Cowan for sharing essential materials with us during the supply chain shortage. This work was part of the DOE Joint BioEnergy Institute (https://www.jbei.org) supported by the U.S. Department of Energy, Office of Science, Office of Biological and Environmental Research, supported by the U.S. Department of Energy, Energy Efficiency and Renewable Energy, Bioenergy Technologies Office, through contract DE-AC02-05CH11231 between Lawrence Berkeley National Laboratory and the U.S. Department of Energy. A.A.N. was supported by a National Science Foundation Graduate Research Fellowship, fellow ID [2018253421]. The views and opinions of the authors expressed herein do not necessarily state or reflect those of the United States Government or any agency thereof. Neither the United States Government nor any agency thereof, nor any of their employees, makes any warranty, expressed or implied, or assumes any legal liability or responsibility for the accuracy, completeness, or usefulness of any information, apparatus, product, or process disclosed, or represents that its use would not infringe privately owned rights. The United States Government retains and the publisher, by accepting the article for publication, acknowledges that the United States Government retains a nonexclusive, paid-up, irrevocable, worldwide license to publish or reproduce the published form of this manuscript, or allow others to do so, for United States Government purposes. The Department of Energy will provide public access to these results of federally sponsored research in accordance with the DOE Public Access Plan (http://energy.gov/downloads/doe-public-access-plan).

