## Supplemental materials for "Characterization and diversification of AraC/XylS family regulators guided by transposon sequencing"

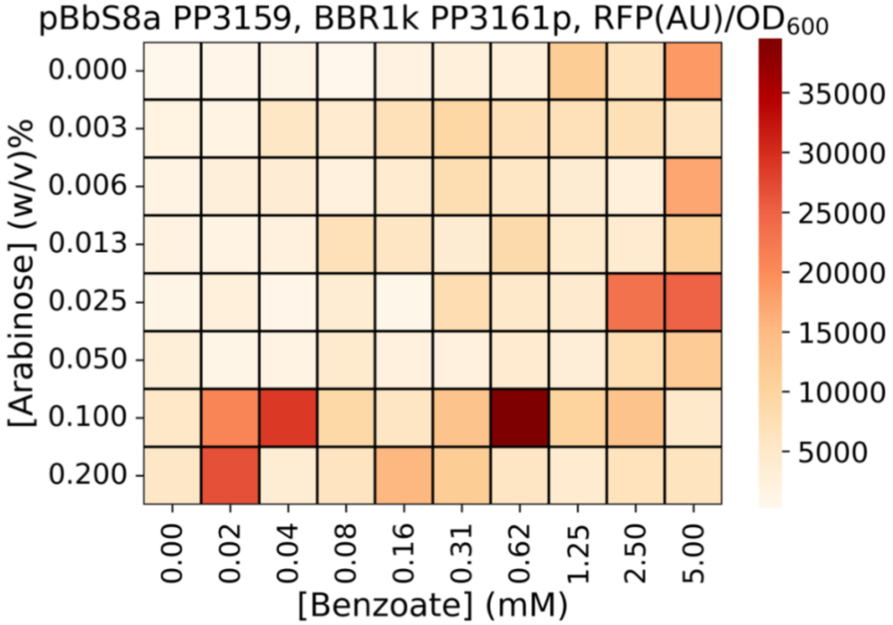


Figure S1: Standard deviation of BenR two plasmid assay shown in Figure 3B (n=3).

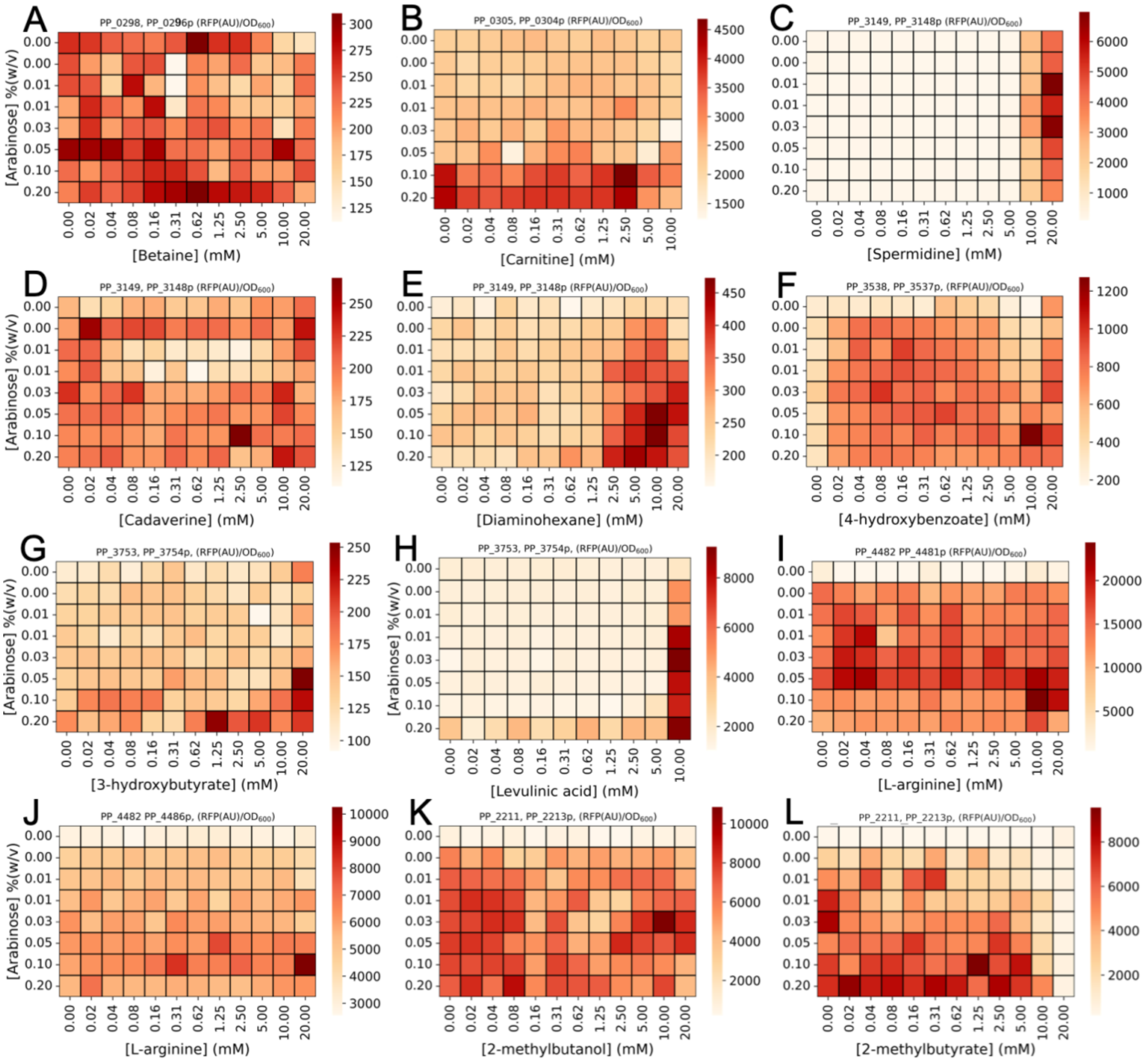


Figure S2: Two-plasmid assays in *E. coli*. Arabinose concentration is varied on the y-axis and the concentration of the potential inducers is varied on the x-axis. Systems and inducers are as follows: A) PP_0298 + $P_{PP\_0296}$ (sIP13) with betaine. B) PP_00305 + $P_{PP\_0304}$ (sIP14) with carnitine . C) PP_3149 + $P_{PP\_3148}$ (sIP15) with spermidine. D) PP_3149 + $P_{PP\_3148}$ (sIP15) with cadaverine. E) PP_3149 + $P_{PP\_3148}$ (sIP15) with 1,6 diaminohexane. F) PP_3538 + $P_{PP\_3537}$ (sIP18) with 4-hydroxybenzoate. G) PP_3753 + $P_{PP\_3754}$ (sIP19) with 3-hydroxybutyrate. H) PP_3753 + $P_{PP\_3754}$ (sIP19) with levulinic acid. I) PP_4482 + $P_{PP\_4481}$ (sIP21) with L-arginine. J) PP_4482 + $P_{PP\_4486}$ (sIP20) with L-arginine. K) PP_2211 + $P_{PP\_2213}$ (sIP23) with 2-methylbutanol. L) PP_2211 + $P_{PP\_2213}$ (sIP23) with 2-methylbutyrate.

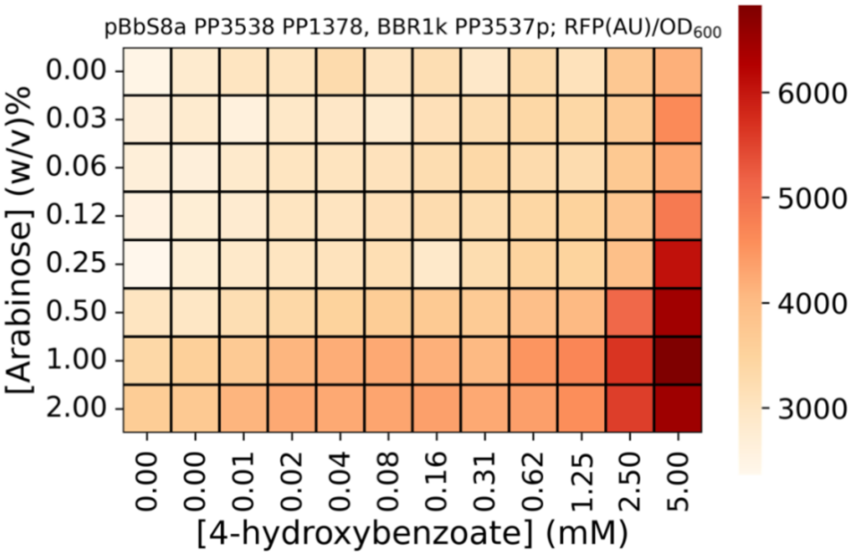


Figure S3: Characterization of a strain (sIP26) carrying aPobR one plasmid system (pIP36) and the transporter PP1378 under the AraC/$P_{BAD}$ system (pIP11). Cultures were grown in LB for 24 hours.

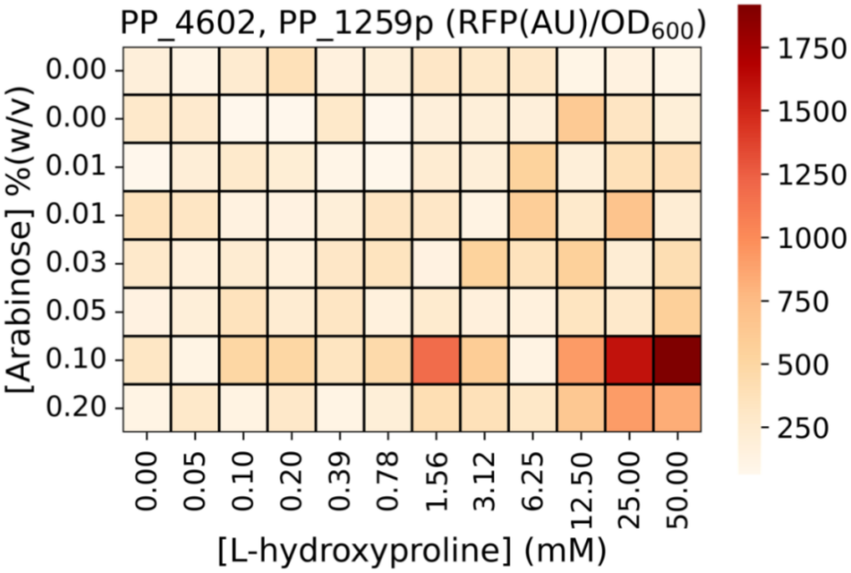


Figure S4: Standard deviation of lhpR two plasmid assay shown in Figure 5A (n=3).

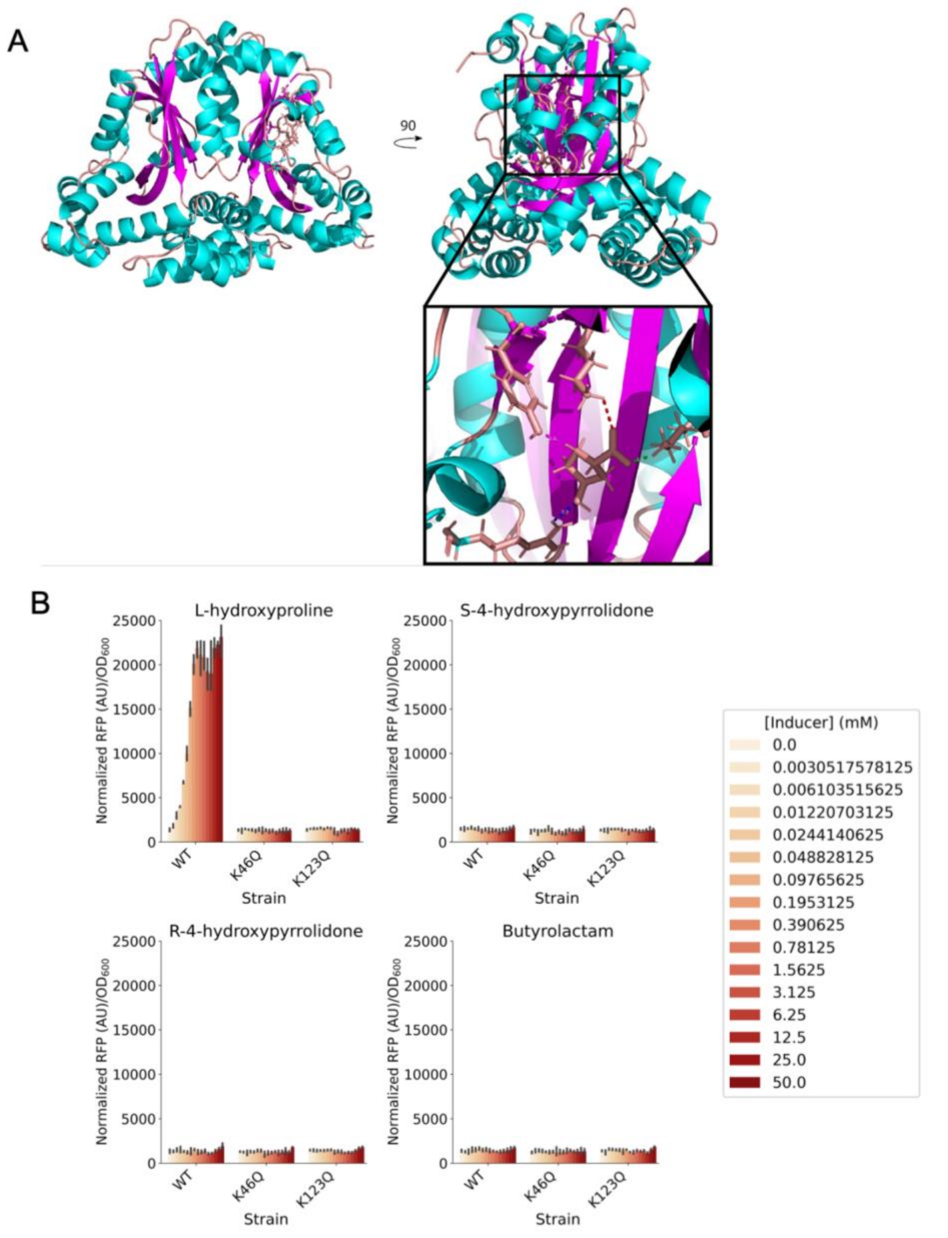


Figure S5: A) Protein structure prediction of PP_4602 (LhpR) as a dimer. The structure was determined with the Foldy implementation of AlphaFold, repacked with sidechain and backbone energy minimization in Foldit-Standalone, and L-HPro docked with SwissDock. K46, K123, R62, and Y52 are depicted in the bottom right panel forming H-bonds with L-HPro. B) RFP measurements in the presence of L-hydroxyproline, butyrolactam, and R/S 4-hydroxypyrrolidone for PP_4602 mutants (pIP40-pIP41, sIP46-sIP47) in the pBBR1k p1_PP4602 PP1259, $P_{PP\_1259}$ RFP system (pIP27) (n = 3, error bars = 95% confidence interval).

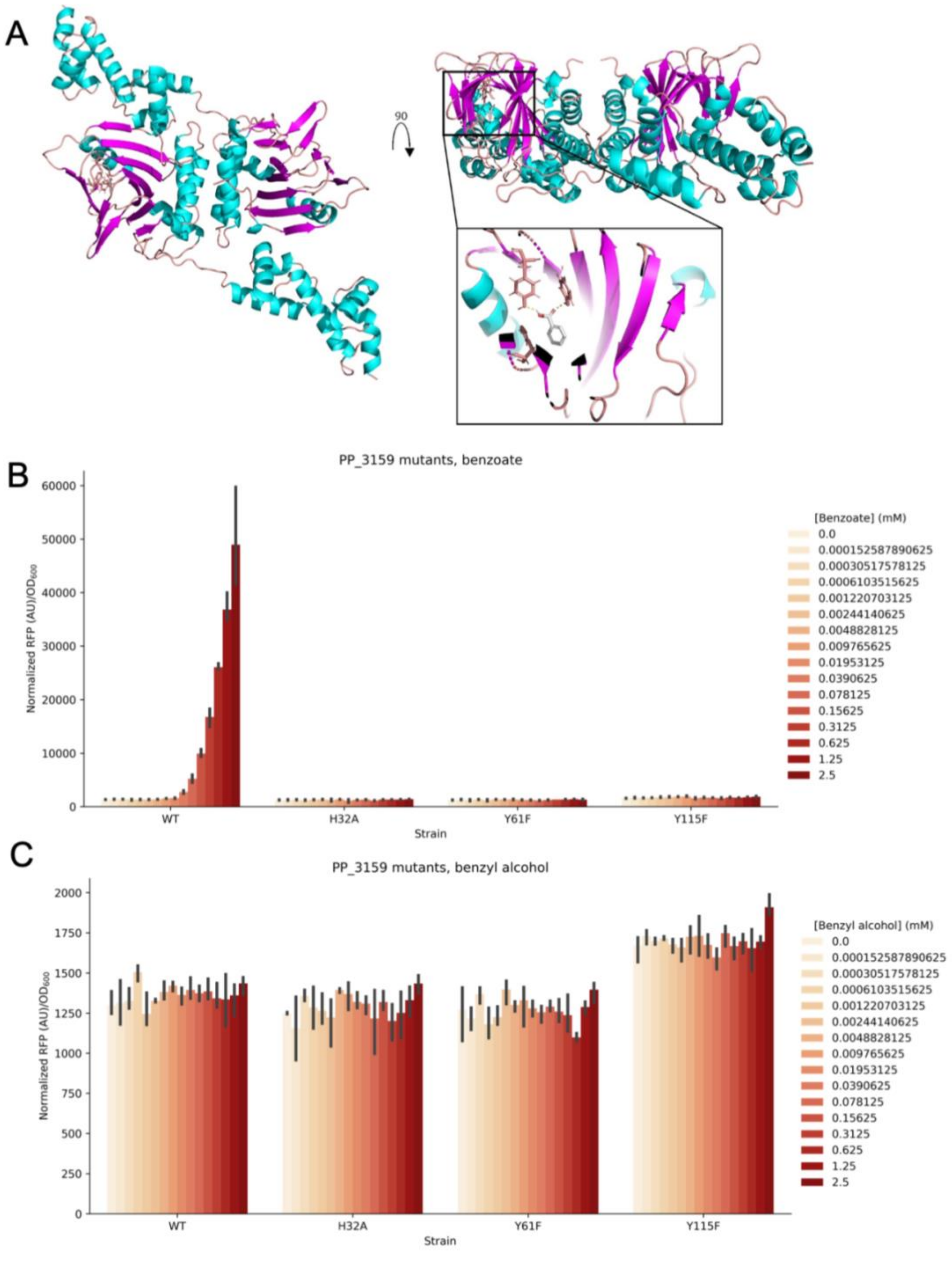


Figure S6: A) AlphaFold predicted structure of BenR docked to benzoate with AutoDock Vina using the Foldy UI. H32, Y61, and Y115 are depicted in the bottom right panel. B) RFP measurements in the presence of benzoate and benzyl alcohol for PP_3159 mutants (pIP37-pIP39, sIP43-sIP45) in the BBR1k p1_PP3159 $P_{PP3161}$ RFP (pIP33) system (n = 3, error bars = 95% confidence interval).

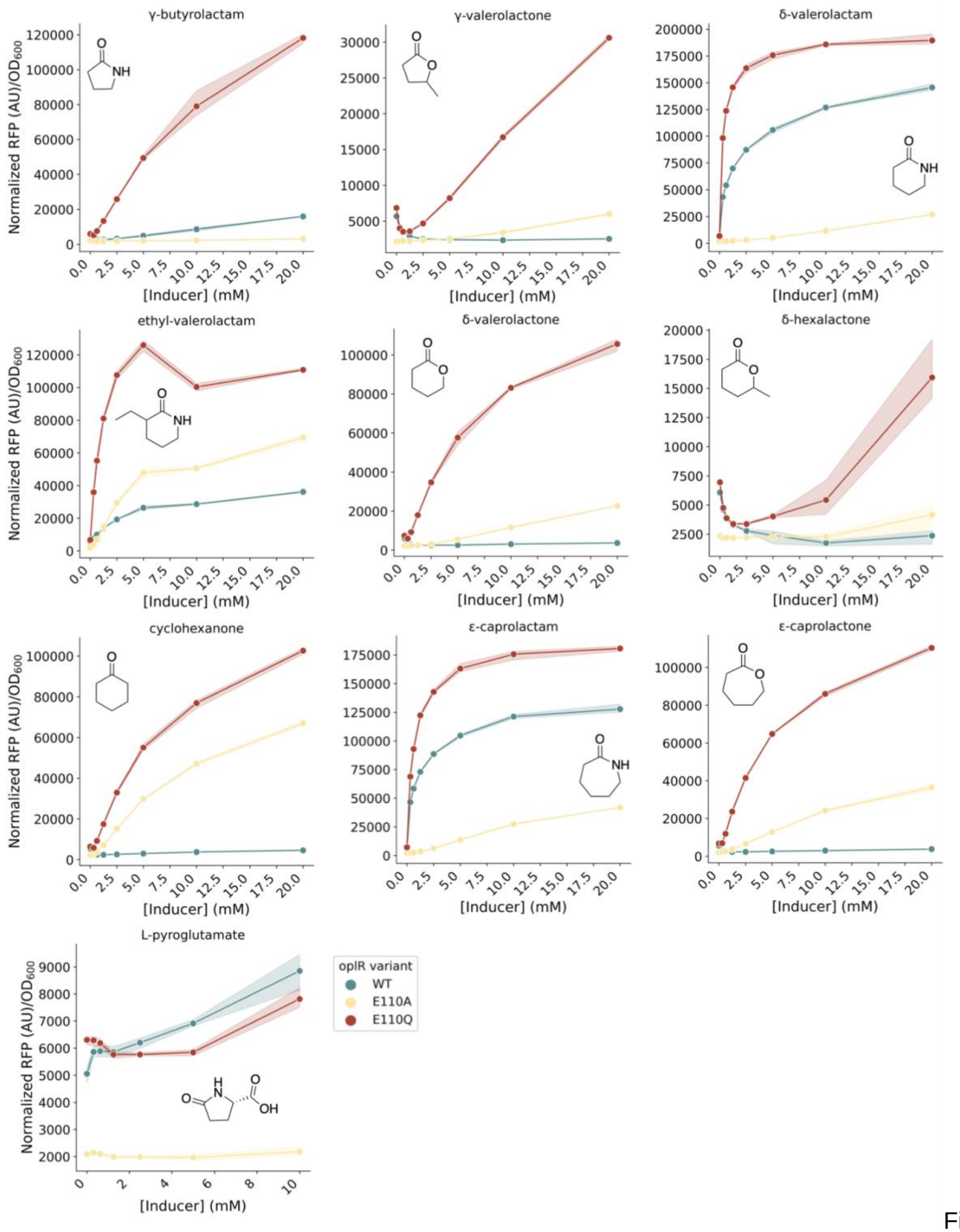


Figure S7: Alternative view of the data presented in Figure 8. Response of oplR wild-type and mutant variants (sIP48-sIP49) to ten cyclic inducers (n = 3, error bars = 95% confidence interval).

Table S1: Plasmids used in this work.

| Abbre-viated name | Plasmid | Description | Source or reference | JBEI part ID |
|---|---|---|---|---|
| | pBbS8a | pSC101 origin with carbenicillin resistance and arabinose inducible promoter | [55] | |
| pIP1 | pBbS8a PP0298 | pBbS8a expressing the ATF PP_0298 | This work | JPUB_021355 |
| pIP2 | pBbS8a PP0305 | pBbS8a expressing the ATF PP_0305 | This work | JPUB_021358 |
| pIP3 | pBbS8a PP3149 | pBbS8a expressing the ATF PP_3149 | This work | JPUB_021359 |
| pIP4 | pBbS8a PP3159 | pBbS8a expressing the ATF PP_3159 | This work | JPUB_021360 |
| pIP5 | pBbS8a PP3538 | pBbS8a expressing the ATF PP_3538 | This work | JPUB_021361 |
| pIP6 | pBbS8a PP3753 | pBbS8a expressing the ATF PP_3753 | This work | JPUB_021362 |
| pIP7 | pBbS8a PP4482 | pBbS8a expressing the ATF PP_4482 | This work | JPUB_021363 |
| pIP8 | pBbS8a PP2211 | pBbS8a expressing the ATF PP_2211 | This work | JPUB_021364 |
| pIP9 | pBbS8a PP4602 | pBbS8a expressing the ATF PP_4602 | This work | JPUB_021365 |
| pIP10 | pBbS8a PP2213 | pBbS8a expressing the CoA ligase PP_2213 | This work | JPUB_021356 |
| pIP11 | pBbS8a PP1378 | pBbS8a expressing the transporter PP_1378 | This work | JPUB_021366 |
| | pBBR1k | pBADT derived broad host-range expression vector with araC and PBAD removed | [23] | |
| pIP12 | pBBR1k PP0296p RFP | Reporter vector with region 200 bp upstream of PP_0296 used as promoter for RFP | This work | JPUB_021368 |
| pIP13 | pBBR1k PP0304p RFP | Reporter vector with region 200 bp upstream of PP_0304 used as promoter for RFP | This work | JPUB_021369 |
| pIP14 | pBBR1k PP3148p RFP | Reporter vector with region 200 bp upstream of PP_3148 used as promoter for RFP | This work | JPUB_021370 |
| pIP15 | pBBR1k PP3161p RFP | Reporter vector with region 200 bp upstream of PP_3161 used as promoter for RFP | This work | JPUB_021371 |
| pIP16 | pBBR1k PP3160p RFP | Reporter vector with region 200 bp upstream of PP_3160 used as promoter for RFP | This work | JPUB_021372 |
| pIP17 | pBBR1k PP3537p RFP | Reporter vector with region 200 bp upstream of PP_3537 used as promoter for RFP | This work | JPUB_021373 |
| pIP18 | pBBR1k PP3754p RFP | Reporter vector with region 200 bp upstream of PP_3754 used as promoter for RFP | This work | JPUB_021374 |
| pIP19 | pBBR1k PP4486p RFP | Reporter vector with region 200 bp upstream | This work | JPUB_021375 |

| | | | | |
|---|---|---|---|---|
| | | of PP_4486 used as promoter for RFP | | |
| pIP20 | pBBR1k PP4481p RFP | Reporter vector with region 200 bp upstream of PP_4481 used as promoter for RFP | This work | JPUB_021376 |
| pIP21 | pBBR1k PP2212p RFP | Reporter vector with region 200 bp upstream of PP_2212 used as promoter for RFP | This work | JPUB_021377 |
| pIP22 | pBBR1k PP2213p RFP | Reporter vector with region 200 bp upstream of PP_2213 used as promoter for RFP | This work | JPUB_021378 |
| pIP23 | pBBR1k PP1259p RFP | Reporter vector with region 200 bp upstream of PP_1259 used as promoter for RFP | This work | JPUB_021379 |
| pIP24 | pBBR1k p1_PP4602, PP1259p RFP | Single plasmid sensor system with PP_4602 under J23103 promoter, and RFP under control of $P_{PP\_1259}$ | This work | JPUB_021380 |
| pIP25 | pBBR1k p2_PP4602, PP1259p RFP | Single plasmid sensor system with PP_4602 under J23114 promoter, and RFP under control of $P_{PP\_1259}$ | This work | JPUB_021382 |
| pIP26 | pBBR1k p3_PP4602, PP1259p RFP | Single plasmid sensor system with PP_4602 under J23105 promoter, and RFP under control of $P_{PP\_1259}$ | This work | JPUB_021384 |
| pIP27 | pBBR1k p1_PP4602 PP1259, PP1259p RFP | Single plasmid sensor system with PP_4602 and PP_1259 under J23103 promoter, and RFP under control of $P_{PP\_1259}$ | This work | JPUB_021386 |
| pIP28 | pBBR1k p2_PP4602 PP1259, PP1259p RFP | Single plasmid sensor system with PP_4602 and PP_1259 under J23114 promoter, and RFP under control of $P_{PP\_1259}$ | This work | JPUB_021388 |
| pIP29 | pBBR1k p3_PP4602 PP1259, PP1259p RFP | Single plasmid sensor system with PP_4602 and PP_1259 under J23105 promoter, and RFP under control of $P_{PP\_1259}$ | This work | JPUB_021390 |
| pIP30 | pBBR1k p1_PP2211, PP2213p RFP | Single plasmid sensor system with PP_2211 under J23103 promoter, and RFP under control of $P_{PP\_2213}$ | This work | JPUB_021398 |
| pIP31 | pBBR1k p2_PP2211, PP2213p RFP | Single plasmid sensor system with PP_2211 under J23114 promoter, and RFP under control of $P_{PP\_2213}$ | This work | JPUB_021400 |
| pIP32 | pBBR1k p3_PP2211, PP2213p RFP | Single plasmid sensor system with PP_2211 under J23105 promoter, and RFP under control of $P_{PP\_2213}$ | This work | JPUB_021402 |
| pIP33 | pBBR1k p1_PP3159, PP3161p RFP | Single plasmid sensor system with PP_3159 under J23103 promoter, and RFP under control of $P_{PP\_3161}$ | This work | JPUB_021392 |

| | | | | |
|---|---|---|---|---|
| pIP34 | pBBR1k p2_PP3159, PP3161p RFP | Single plasmid sensor system with PP_3159 under J23114 promoter, and RFP under control of $P_{PP\_3161}$ | This work | JPUB_021394 |
| pIP35 | pBBR1k p3_PP3159, PP3161p RFP | Single plasmid sensor system with PP_3159 under J23105 promoter, and RFP under control of $P_{PP\_3161}$ | This work | JPUB_021397 |
| pIP36 | BBR1k_p2_PP3538 PP3537p RFP | Single plasmid sensor system with PP_3538 under J23105 promoter, and RFP under control of $P_{PP\_3537}$ | This work | JPUB_021421 |
| pIP37 | pBBR1k p1_PP3159_H32A PP3161p RFP | Single plasmid sensor system with PP_3159_H32A under J23103 promoter, and RFP under control of $P_{PP\_3161}$ | This work | JPUB_021408 |
| pIP38 | pBBR1k p1_PP3159_Y61F PP3161p RFP | Single plasmid sensor system with PP_3159_Y61F under J23103 promoter, and RFP under control of $P_{PP\_3161}$ | This work | JPUB_021410 |
| pIP39 | pBBR1k p1_PP3159_Y115F PP3161p RFP | Single plasmid sensor system with PP_3159_Y115F under J23103 promoter, and RFP under control of $P_{PP\_3161}$ | This work | JPUB_021412 |
| pIP40 | pBBR1k p1_PP4602_K123Q PP1259, PP1259p RFP | Single plasmid sensor system with PP_4602_K123Q and PP_1259 under J23103 promoter, and RFP under control of $P_{PP\_1259}$ | This work | JPUB_021414 |
| pIP41 | pBBR1k p1_PP4602_K46Q PP1259, PP1259p RFP | Single plasmid sensor system with PP_4602_K46Q and PP1259 under J23103 promoter, and RFP under control of $P_{PP\_1259}$ | This work | JPUB_021416 |
| | pLacSens1 | Lactam biosensor based on the AFR OplR, BBR1k derived backbone | [23] | |
| pIP42 | pLacSens1 E110A | E110A mutation of oplR in pLacSens1 | This work | JPUB_021404 |
| pIP43 | pLacSens1 E110Q | E110Q mutation of oplR in pLacSens1 | This work | JPUB_021406 |
| | pMQ30 | Suicide vector for allelic replacement with $Gm^r$, SacB | | |
| pIP44 | pMQ30 ΔPP2211 | Suicide vector for in-frame delection of PP2211 | This work | JPUB_021422 |
| pIP45 | pMQ30 ΔPP2213 | Suicide vector for in-frame delection of PP2213 | This work | JPUB_021424 |
| pIP46 | pMQ30 ΔPP2216 | Suicide vector for in-frame delection of PP2216 | This work | JPUB_021426 |

Table S2: Strains used in this work.

| Abbre-viated name | Name | Description | Source or reference | JBEI part ID |
|---|---|---|---|---|
| | *E. coli* XL1-blue | | Agilent | |
| | *Pseudomonas putida* KT2440 | | ATCC 47054 | |
| | *E. coli* S17 | | ATCC 47055 | |
| sIP1 | *P. putida* pBBR1k PP0296p RFP | *P. putida* carrying the reporter plasmid pBBR1k PP0296p RFP (pIP12) | This work | JPUB_021325 |
| sIP2 | *P. putida* pBBR1k PP0304p RFP | *P. putida* carrying the reporter plasmid pBBR1k PP0304p RFP (pIP13) | This work | JPUB_021309 |
| sIP3 | *P. putida* pBBR1k PP3148p RFP | *P. putida* carrying the reporter plasmid pBBR1k PP3148p RFP (pIP14) | This work | JPUB_021311 |
| sIP4 | *P. putida* pBBR1k PP3161p RFP | *P. putida* carrying the reporter plasmid pBBR1k PP3161p RFP (pIP15) | This work | JPUB_021313 |
| sIP5 | *P. putida* pBBR1k PP3160p RFP | *P. putida* carrying the reporter plasmid pBBR1k PP3160p RFP (pIP16) | This work | JPUB_021315 |
| sIP6 | *P. putida* pBBR1k PP3537p RFP | *P. putida* carrying the reporter plasmid pBBR1k PP3537p RFP (pIP17) | This work | JPUB_021317 |
| sIP7 | *P. putida* pBBR1k PP3754p RFP | *P. putida* carrying the reporter plasmid pBBR1k PP3754p RFP (pIP18) | This work | JPUB_021319 |
| sIP8 | *P. putida* pBBR1k PP4486p RFP | *P. putida* carrying the reporter plasmid pBBR1k PP4486p RFP (pIP19) | This work | JPUB_021321 |
| sIP9 | *P. putida* pBBR1k PP4481p RFP | *P. putida* carrying the reporter plasmid pBBR1k PP4481p RFP (pIP20) | This work | JPUB_021323 |
| sIP10 | *P. putida* pBBR1k PP2212p RFP | *P. putida* carrying the reporter plasmid pBBR1k PP2212p RFP (pIP21) | This work | JPUB_021331 |
| sIP11 | *P. putida* pBBR1k PP2213p RFP | *P. putida* carrying the reporter plasmid pBBR1k PP2213p RFP (pIP22) | This work | JPUB_021333 |
| sIP12 | *P. putida* pBBR1k PP1259p RFP | *P. putida* carrying the reporter plasmid pBBR1k PP1259p RFP (pIP23) | This work | JPUB_021334 |
| sIP13 | PP0298, PP0296p RFP | Two plasmid sensor comprised of pBbS8a PP0298 (pIP1) and pBBR1k PP0296p RFP (pIP12) in *E. coli* XL1-Blue | This work | JPUB_021335 |
| sIP14 | PP0305, PP0304p RFP | Two plasmid sensor comprised of pBbS8a PP0305 (pIP2) and pBBR1k PP0304p RFP (pIP13) in *E. coli* XL1-Blue | This work | JPUB_021337 |
| sIP15 | PP3149, PP3148p RFP | Two plasmid sensor comprised of pBbS8a PP3149 (pIP3) and pBBR1k PP3148p RFP | This work | JPUB_021339 |

| | | | | |
|---|---|---|---|---|
| | | (pIP14) in *E. coli* XL1-Blue | | |
| sIP16 | PP3159, PP3161p RFP | Two plasmid sensor comprised of pBbS8a PP3159 (pIP4) and pBBR1k PP3161p RFP (pIP15) in *E. coli* XL1-Blue | This work | JPUB_021341 |
| sIP17 | PP3159, PP3160p RFP | Two plasmid sensor comprised of pBbS8a PP3159 (pIP4) and pBBR1k PP3160p RFP (pIP16) in *E. coli* XL1-Blue | This work | JPUB_021343 |
| sIP18 | PP3538, PP3537p RFP | Two plasmid sensor comprised of pBbS8a PP3538 (pIP5) and pBBR1k PP3537p RFP (pIP17) in *E. coli* XL1-Blue | This work | JPUB_021344 |
| sIP19 | PP3753, PP3754p RFP | Two plasmid sensor comprised of pBbS8a PP3753 (pIP6) and pBBR1k PP3754p RFP (pIP18) in *E. coli* XL1-Blue | This work | JPUB_021346 |
| sIP20 | PP4482, PP4486p RFP | Two plasmid sensor comprised of pBbS8a PP4482 (pIP7) and pBBR1k PP4486p RFP (pIP19) in *E. coli* XL1-Blue | This work | JPUB_021348 |
| sIP21 | PP4482, PP4481p RFP | Two plasmid sensor comprised of pBbS8a PP4482 (pIP7) and pBBR1k PP4481p RFP (pIP19) in *E. coli* XL1-Blue | This work | JPUB_021350 |
| sIP22 | PP2211, PP2212p RFP | Two plasmid sensor comprised of pBbS8a PP2211 (pIP8) and pBBR1k PP2212p RFP (pIP21) in *E. coli* XL1-Blue | This work | JPUB_021351 |
| sIP23 | PP2211, PP2213p RFP | Two plasmid sensor comprised of pBbS8a PP2211 (pIP8) and pBBR1k PP2213p (pIP22) RFP in *E. coli* XL1-Blue | This work | JPUB_021353 |
| sIP24 | PP4602, PP1259p RFP | Two plasmid sensor comprised of pBbS8a PP4602 (pIP9) and pBBR1k PP1259p RFP (pIP23) in *E. coli* XL1-Blue | This work | JPUB_021354 |
| sIP25 | pBbS8a PP2213, pBBR1k p1_PP2211 PP2213p RFP | Two plasmid system comprised of pBbS8a PP2213 (pIP10) and one plasmid system pBBR1k p1_PP2211 PP2213p RFP (pIP30) in *E. coli* XL1-Blue | This work | JPUB_021418 |
| sIP26 | pBbS8a PP1378, pBBR1k p2_ PP3538 PP3537p RFP | Two plasmid system comprised of pBbS8a PP1378 (pIP11) and pBBR1k p2_ PP3538 PP3537p_RFP (pIP36) in *E. coli* XL1-Blue | This work | JPUB_021419 |
| sIP27 | ΔPP2211 pBBR1k PP2213p RFP | *P. putida* deletion strain of the ATF PP_2211 carrying the pBBR1k PP2213p RFP (pIP22) reporter plasmid | This work | JPUB_021329 |
| sIP28 | ΔPP2213 pBBR1k PP2213p RFP | *P. putida* deletion strain of the CoA ligase PP_2213 carrying the pBBR1k PP2213p RFP (pIP22) reporter plasmid | This work | JPUB_021330 |
| sIP29 | ΔPP2216 pBBR1k PP2213p RFP | *P. putida* deletion strain of the acyl CoA dehydrogenase PP_2216 carrying the pBBR1k PP2213p RFP (pIP22) reporter plasmid | This work | JPUB_021327 |

| | | | | |
|---|---|---|---|---|
| sIP30 | pBBR1k p1_PP4602, PP1259p RFP | *E. coli* XL1-Blue carrying the pBBR1k p1_PP4602, PP1259p RFP (pIP24) plasmid | This work | JPUB_021380 |
| sIP31 | pBBR1k p2_PP4602, PP1259p RFP | *E. coli* XL1-Blue carrying the pBBR1k p2_PP4602, PP1259p RFP (pIP25) plasmid | This work | JPUB_021382 |
| sIP32 | pBBR1k p3_PP4602, PP1259p RFP | *E. coli* XL1-Blue carrying the pBBR1k p3_PP4602, PP1259p RFP (pIP26) plasmid | This work | JPUB_021384 |
| sIP33 | pBBR1k p1_PP4602 PP1259, PP1259p RFP | *E. coli* XL1-Blue carrying the pBBR1k p1_PP4602 PP1259, PP1259p RFP (pIP27) plasmid | This work | JPUB_021386 |
| sIP34 | pBBR1k p2_PP4602 PP1259, PP1259p RFP | *E. coli* XL1-Blue carrying the pBBR1k p2_PP4602 PP1259, PP1259p RFP (pIP28) plasmid | This work | JPUB_021388 |
| sIP35 | pBBR1k p3_PP4602 PP1259, PP1259p RFP | *E. coli* XL1-Blue carrying the pBBR1k p3_PP4602 PP1259, PP1259p RFP (pIP27) plasmid | This work | JPUB_021390 |
| sIP36 | pBBR1k p1_PP2211, PP2213p RFP | *E. coli* XL1-Blue carrying the pBBR1k p1_PP2211, PP2213p RFP (pIP30) plasmid | This work | JPUB_021398 |
| sIP37 | pBBR1k p2_PP2211, PP2213p RFP | *E. coli* XL1-Blue carrying the pBBR1k p2_PP2211, PP2213p RFP (pIP31) plasmid | This work | JPUB_021400 |
| sIP38 | pBBR1k p3_PP2211, PP2213p RFP | *E. coli* XL1-Blue carrying the pBBR1k p3_PP2211, PP2213p RFP (pIP32) plasmid | This work | JPUB_021402 |
| sIP39 | pBBR1k p1_PP3159, PP3161p RFP | *E. coli* XL1-Blue carrying the pBBR1k p1_PP3159, PP3161p RFP (pIP33) plasmid | This work | JPUB_021392 |
| sIP40 | pBBR1k p2_PP3159, PP3161p RFP | *E. coli* XL1-Blue carrying the pBBR1k p2_PP3159, PP3161p RFP (pIP34) plasmid | This work | JPUB_021394 |
| sIP41 | pBBR1k p3_PP3159, PP3161p RFP | *E. coli* XL1-Blue carrying the pBBR1k p3_PP3159, PP3161p RFP (pIP35) plasmid | This work | JPUB_021397 |
| sIP42 | BBR1k_p2_PP3538 PP3537p RFP | *E. coli* XL1-Blue carrying the BBR1k_p2_PP3538 PP3537p RFP (pIP36) plasmid | This work | JPUB_021421 |
| sIP43 | pBBR1k p1_PP3159_H32A PP3161p RFP | *E. coli* XL1-Blue carrying the pBBR1k p1_PP3159_H32A PP3161p RFP (pIP37) plasmid | This work | JPUB_021408 |
| sIP44 | pBBR1k p1_PP3159_Y61F | *E. coli* XL1-Blue carrying the pBBR1k p1_PP3159_Y61F PP3161p RFP (pIP38) | This work | JPUB_021410 |

| | | | | |
|---|---|---|---|---|
| | PP3161p RFP | plasmid | | |
| sIP45 | pBBR1k p1_PP3159_Y115F PP3161p RFP | *E. coli* XL1-Blue carrying the pBBR1k p1_PP3159_Y115F PP3161p RFP (pIP39) plasmid | This work | JPUB_021412 |
| sIP46 | pBBR1k p1_PP4602_K123Q PP1259, PP1259p RFP | *E. coli* XL1-Blue carrying the pBBR1k p1_PP4602_K123Q PP1259, PP1259p RFP (pIP40) plasmid | This work | JPUB_021414 |
| sIP47 | pBBR1k p1_PP4602_K46Q PP1259, PP1259p RFP | *E. coli* XL1-Blue carrying the pBBR1k p1_PP4602_K46Q PP1259, PP1259p RFP (pIP41) plasmid | This work | JPUB_021416 |
| sIP48 | pLacSens1 E110A | *E. coli* XL1-Blue carrying the pLacSens1 E110A (pIP42) plasmid | This work | JPUB_021404 |
| sIP49 | pLacSens1 E110Q | *E. coli* XL1-Blue carrying the pLacSens1 E110A (pIP43) plasmid | This work | JPUB_021406 |

Table S3: Links to public Foldy predicted structures and docking for AFRs in this work.

| **Predicted AFR** | **Link to Foldy public structure** |
|---|---|
| PP_0298 (gdbR) | Monomer: https://foldy.lbl.gov/fold/793<br>Dimer: https://foldy.lbl.gov/fold/2483 |
| PP_0305 (cdhR) | Monomer: https://foldy.lbl.gov/fold/798<br>Dimer: https://foldy.lbl.gov/fold/2481 |
| PP_3149 | Monomer: https://foldy.lbl.gov/fold/1253<br>Dimer: https://foldy.lbl.gov/fold/2484 |
| PP_3538 (pobR) | Monomer: https://foldy.lbl.gov/fold/1313<br>Dimer: https://foldy.lbl.gov/fold/2526 |
| PP_3753 | Monomer: https://foldy.lbl.gov/fold/1360<br>Dimer: https://foldy.lbl.gov/fold/2532 |
| PP_4602 | Monomer: https://foldy.lbl.gov/fold/1794<br>Truncated monomer: https://foldy.lbl.gov/fold/1795<br>Dimer: https://foldy.lbl.gov/fold/1793<br>Truncated dimer: https://foldy.lbl.gov/fold/1796<br>Ligand binding domain: https://foldy.lbl.gov/fold/2541 |
| PP_3159 (benR) | Monomer: https://foldy.lbl.gov/fold/1255<br>Dimer: https://foldy.lbl.gov/fold/2528 |
| PP_4482 | Monomer: https://foldy.lbl.gov/fold/1493<br>Dimer: https://foldy.lbl.gov/fold/2482 |
| PP_2211 (tmbR) | Monomer: https://foldy.lbl.gov/fold/1149<br>Dimer: https://foldy.lbl.gov/fold/2487 |
| PP_3516 (oplR) | Monomer: https://foldy.lbl.gov/fold/1310<br>Dimer: https://foldy.lbl.gov/fold/2449 |
